# Common γ-chain cytokines induce an epigenomically plastic precursor-like KIT^+^ ILC2 state linked to immune disease susceptibility

**DOI:** 10.64898/2026.02.27.708582

**Authors:** Simone E. M. Olsthoorn, Anne Onrust-Van Schoonhoven, Marjolein J.W. de Bruijn, Menno van Nimwegen, Gregory van Beek, Willem de Koning, Lianne Trap, Esmee K. van der Ploeg, Mathijs A. Sanders, Laura Surace, James P. Di Santo, Rudi W. Hendriks, Ralph Stadhouders

**Affiliations:** Department of Pulmonary Medicine, Erasmus MC University Medical Center, Rotterdam, The Netherlands; Department of Hematology, Erasmus MC University Medical Center, Rotterdam, The Netherlands; Department of Internal Medicine, Erasmus MC University Medical Center, Rotterdam, The Netherlands; Institute of Clinical Chemistry and Clinical Pharmacology, University Hospital Bonn, University of Bonn, Germany; Innate Immunity Unit, Institut Pasteur, Université Paris Cité, Inserm U1223, Paris, France

**Keywords:** ILC2, ILCP, epigenome, plasticity, KIT, cytokine signaling, STAT5

## Abstract

**Background:** Group 2 innate lymphoid cells (ILC2s) are key effector cells of type-2 immunity. A subset of ILC2s expresses KIT (CD117), which display increased phenotypic plasticity and were previously linked to severe asthma and psoriasis. However, the molecular mechanisms promoting a KIT^+^ ILC2 state remain poorly understood.

**Objective:** Define the molecular basis for the enhanced plasticity of KIT^+^ ILC2s and identify signals that induce this phenotype, including links with immune disease susceptibility.

**Methods:** We combine bulk as well as single-cell transcriptome (RNA-seq) and epigenome (ATAC-seq) with *in vitro* culture assays using primary human KIT^+^ or KIT^neg^ ILC2s and multipotent ILC precursors (ILCPs). Epigenomic data were integrated with genetic risk variants for major human immune diseases.

**Results:** Multi-omics analyses revealed that KIT^+^ ILC2s maintain a unique hybrid character marked by expression and open chromatin of genes linked to both ILCP and ILC2 biology. KIT^+^ ILC2s showed extensive epigenomic priming at gene loci related to naive lymphocyte biology, tissue homing, and ILC3 effector functions, including *IL17* and *IL23R* – explaining why KIT^+^ ILC2s are poised to adopt an ILC3-like phenotype. Genetic risk variants for asthma and autoimmunity are enriched in the poised epigenome of KIT^+^ ILC2s. Common γ-chain cytokines IL-2 and IL-7 induced a KIT^+^ phenotype in KIT^neg^ ILC2s through STAT5 activation.

**Conclusions:** Our study defines KIT^+^ ILC2s as a developmentally immature state carrying a precursor-like epigenome that promotes phenotypic plasticity and is linked to immune disease susceptibility. Importantly, we identify STAT5-mediated cytokine signals as candidates for therapeutic targeting of KIT^+^ ILC2s.

## INTRODUCTION

Innate lymphoid cells (ILCs) play important roles in immune responses against pathogens and in maintaining tissue homeostasis^1–3^. Akin to their adaptive T helper (Th) cell counterparts, ILCs consist of group 1 (ILC1), group 2 (ILC2) and group 3 (ILC3) cells, which in terms of transcription factor (TF) dependencies and cytokine production, mirror Th1, Th2, and Th17 cells, respectively^4,5^. In contrast to T cells, ILCs lack antigen-specific receptors and depend on microenvironmental signals for their activation.

ILC2s are key mediators of protective type-2 immune responses against parasites but also drive pathological inflammation in diseases such as asthma. Dependent on the TF GATA3, ILC2s produce large amounts of interleukin 5 (IL-5) and IL-13 to induce eosinophilic inflammation in response to tissue alarmin signals such as IL-25, IL-33, and thymic stromal lymphopoietin (TSLP)^6–8^. Additionally, common γ-chain cytokines such as IL-2 and IL-7 are essential for ILC2 development and maintenance^6,8^. Although ILC2s are mostly tissue-resident, their activation can induce inter-organ trafficking to disseminate type-2 immunity^9–12^. Although ILC2 development remains poorly understood, mature human ILC2s are thought to derive from unipotent and multipotent ILC progenitors (ILCP) upon cytokine exposure^13,14^. Like ILC2s, ILCPs can be found in various (mucosal) tissues and in the blood^13–16^, in line with local ILCP differentiation to replenish tissue ILC populations^17^. ILCPs express KIT (or CD117), a receptor tyrosine kinase commonly expressed on hematopoietic stem and progenitor cells. A subset of ILC2s also expresses KIT^18,19^, although it is unclear whether these cells represent a more immature ILC2 state.

ILC2s can be highly plastic, allowing them to acquire functional characteristics normally linked to other ILC subsets^20^. Driven by changes in their cytokine environment, human ILC2s can adopt ILC1-like or ILC3-like phenotypes^21–26^. ILC2 plasticity appears especially prevalent in chronic inflammatory diseases of the airways (e.g., cystic fibrosis^24^, COPD, asthma^21,22^) and skin (e.g., psoriasis^25^, atopic dermatitis^27^). KIT^+^ ILC2s – making up over half of all circulating human ILC2s – were recently postulated to be particularly sensitive to cytokine-driven phenotypic plasticity^26^. Specific cytokine milieus induced more potent IFNγ (type-1) or IL-17 (type-3) production in KIT^+^ ILC2s as compared to their KIT^neg^ counterparts^26^. IL-17 expression by KIT^+^ ILC2s was associated with upregulation of markers linked to ILC3 (e.g., the TF RORγt) and concomitant downregulation of GATA3^25,26^. In contrast, KIT^neg^ ILC2s appear less plastic and may represent a more lineage-committed subset^12,25,26^. Importantly, KIT^+^ ILC2s may represent major producers of IL-17 in psoriasis and asthma, acting as drivers of severe (neutrophilic) inflammation^25,28,29^. How KIT^+^ ILC2s maintain a high capacity for phenotypic plasticity and whether this is linked to their role in inflammatory diseases is unknown.

Epigenomes provide information on the developmental history and present identity of cells, but also on their future responsive potential^30–32^. We and others previously postulated that ILC2s maintain a broadly permissive epigenome that primes them for rapid adaptation of their gene expression program to microenvironmental changes^33–35^. However, whether such epigenetic priming also underlies human ILC2 differentiation and plasticity – in particular of the KIT^+^ subset – is unknown. Moreover, the signals that maintain a KIT^+^ state in human ILC2s are poorly understood. As a consequence, precise developmental relationships between KIT^+^ ILC2s, KIT^neg^ ILC2s, and ILCPs remain unclear. Here, we set out to address these questions using multi-omics analyses and *in vitro* culture assays with primary human ILCs.

## METHODS

A detailed description of all methodology can be found in the Supplementary Material.

## RESULTS

### Transcriptional profiling of circulating human KIT^+^ and KIT^neg^ ILC2s

To better understand how KIT^+^ ILC2s relate to both KIT^+^ ILCPs and KIT^neg^ ILC2s at a transcriptomic and epigenomic level, we isolated these three subsets from healthy donor blood samples and measured genome-wide gene expression and chromatin accessibility using bulk RNA-Seq and ATAC-Seq, respectively (**Fig.1A-B**). Principal component analysis (PCA) at the transcriptome level revealed that gene expression programs in KIT^+^ ILC2s closely resembled those from KIT^neg^ ILC2s but were distinct from those in ILCPs (**Fig.1C**). As expected, canonical Th2/ILC2-defining genes, including TFs *RORA* and *BCL11B*, the *GATA3-AS1* long non-coding RNA^36^, and surface markers *KLRB1* (encoding CD161), *KLRG1*, and *PTGDR2* (encoding CRTH2), demonstrated elevated expression levels in both ILC2 subsets compared to ILCPs (**Fig.1D**). In addition, ILCP showed high expression of *KIT, NCR1* (encoding NKp46), *CCR7*, and several genes essential for mouse ILC development, such as *ID2, TOX*, and *TCF7*^4^ (**Fig.1D**). These observed expression patterns align with findings from previous studies^13,14,25^. Notably, KIT^+^ ILC2s often showed intermediate expression levels of cell identity genes, including *GATA3-AS1* and most ILCP-associated markers (**Fig.1D**). Flow cytometry confirmed similar patterns at the protein level for CRTH2 (*PTGDR2*), KLRG1, NCR1, and KIT (**Fig.1E**).

**Figure 1.**
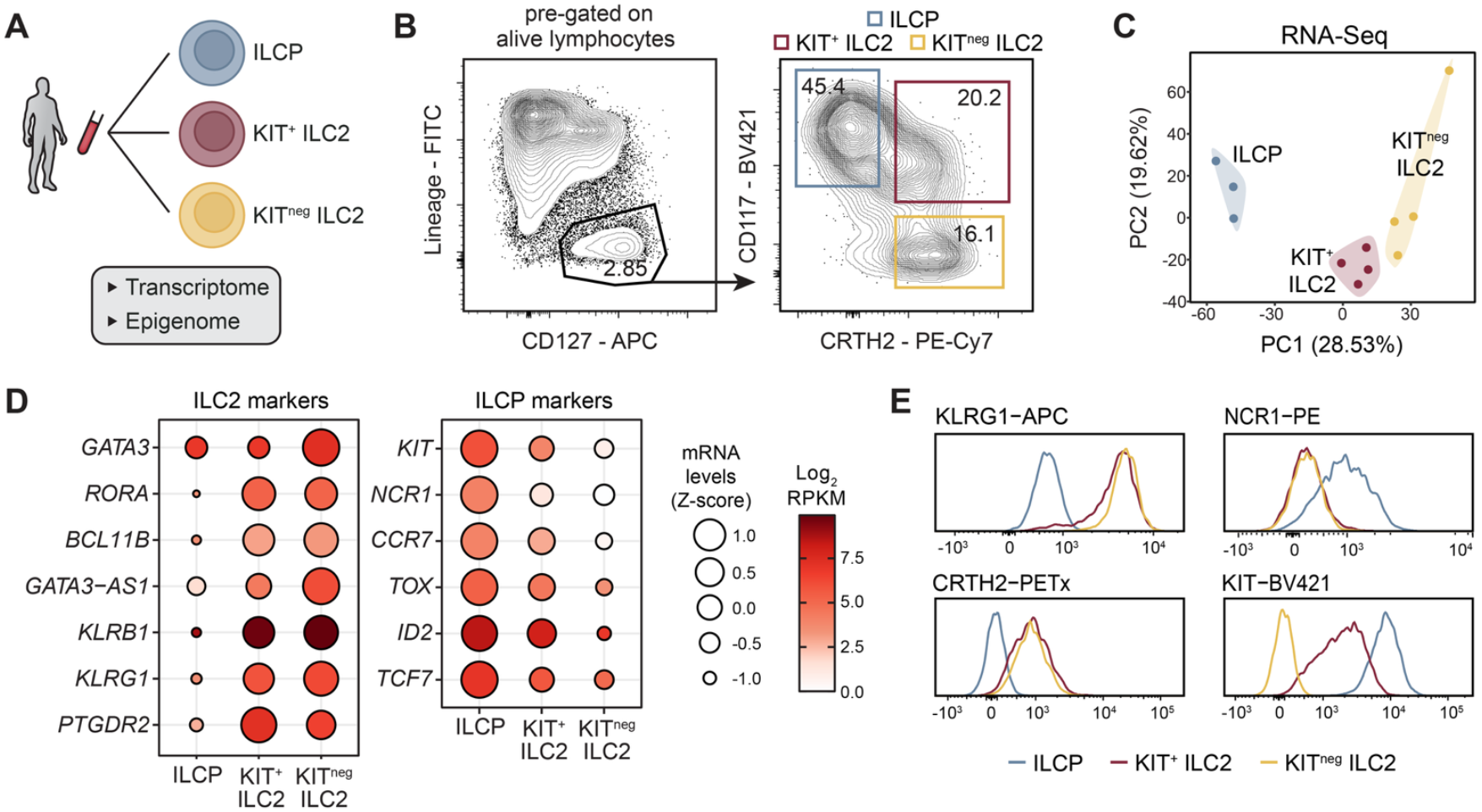
Transcriptional profiling of primary human blood ILCs. (**A**) Experimental setup for conducting transcriptome (RNA-seq) and epigenome (ATAC-seq) analyses of ILCPs, KIT^+^ ILC2s, and KIT^neg^ ILC2s from human peripheral blood. (**B**) Flow cytometry gating strategy to isolate ILCPs, KIT^+^ ILC2s, and KIT^neg^ ILC2s. A representative donor is shown; numbers indicate percentages of gated cells. (**C**) PCA of log_2_-transformed mRNA gene expression values of all genes that were expressed in any of the three ILC subsets (n=3 for ILCP, n=4 for both ILC2 subsets). (**D**) Balloon plot depicting mRNA expression levels of canonical ILC2 and ILCP marker genes in the indicated ILC subsets. Circle size indicates relative expression to the other subsets (row z-score), while the color scale shows absolute expression levels (log_2_ RPKM). (**E**) Flow cytometry measurement of KLRG1, NCR1, CRTH2, and KIT surface protein levels on independently isolated ILCPs, KIT^+^ ILC2s, and KIT^neg^ ILC2s (representative sample shown).

These data show that KIT^+^ ILC2s adopt a global transcriptional program characteristic of the ILC2 lineage, although they show upregulated expression of genes associated with a naive progenitor ILC phenotype.

### KIT^+^ ILC2s share transcriptional programs with both KIT^neg^ ILC2s and ILCPs

Differential gene expression analysis detected fewer differentially expressed genes (DEGs) in KIT^+^ *vs*. KIT^neg^ ILC2s (n=207) than in KIT^+^ ILC2s *vs*. ILCPs (n=945) (**Fig.2A-B**). Over half (52-56%) of the DEGs identified in both ILCP *vs*. ILC2 comparisons were shared (484+42=526 genes; **Fig.2B**).

**Figure 2.**
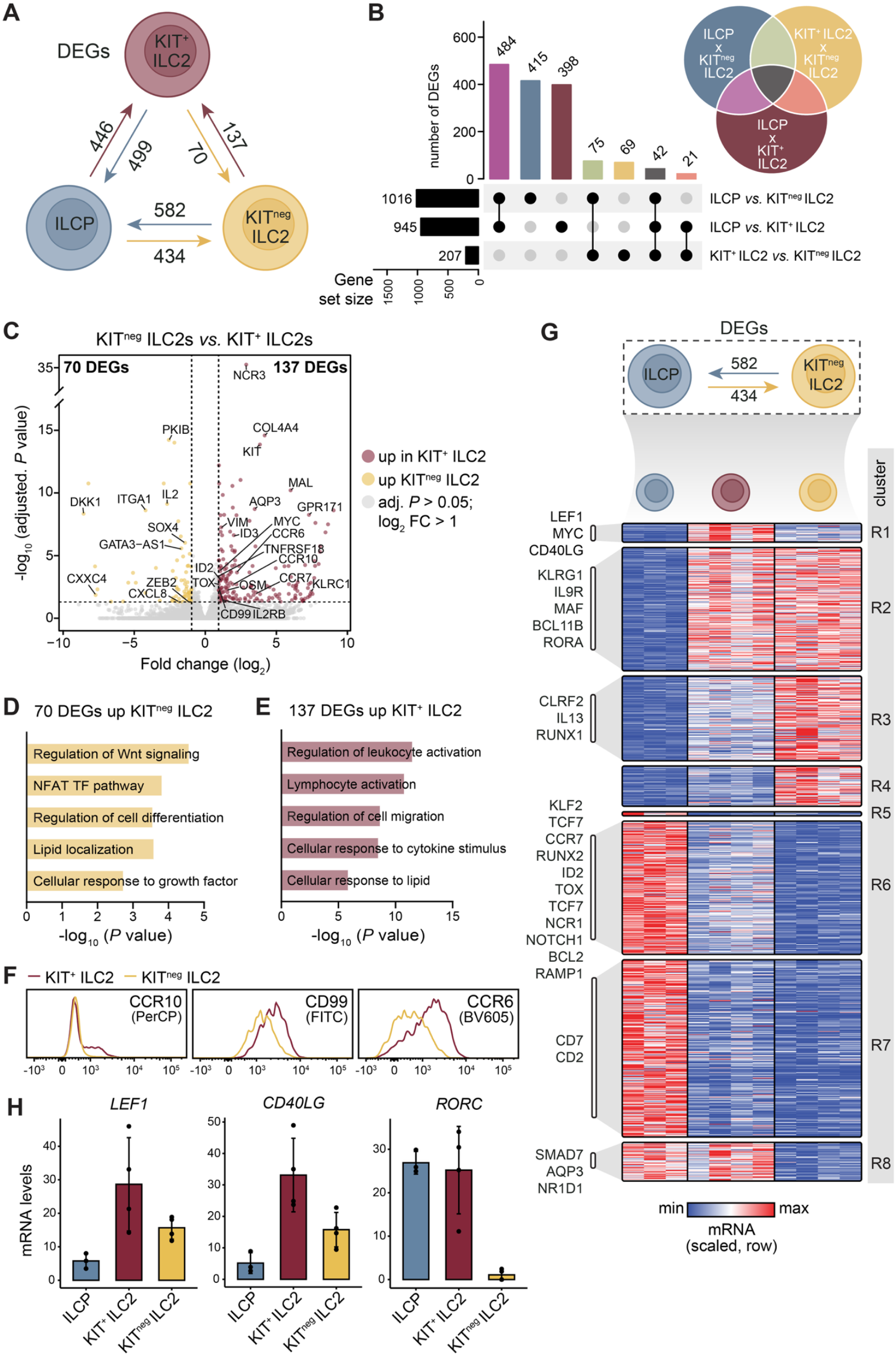
KIT^+^ ILC2s maintain transcriptional programs linked to ILCP biology. (**A**) Schematic overview of differentially expressed genes (DEGs) between ILCPs, KIT^+^ ILC2s, and KIT^neg^ ILC2s (log_2_ fold change > 1, adjusted *P* < 0.05). (**B**) Upset plot visualizing overlapping and unique DEGs among the indicated comparisons of ILC subsets (further illustrated in the Venn diagram; color code matches with bar graph). (**C**) Volcano plot showing DEGs between KIT^+^ ILC2s and KIT^neg^ ILC2s (adjusted P-values shown); selected genes are highlighted. (**D-E**) Bar graphs showing selected significantly enriched pathways for the DEGs between KIT^+^ ILC2s and KIT^neg^ ILC2s. (**F**) Flow cytometry measurement of CCR10, CD99, and CCR6 protein levels on independently isolated KIT^+^ and KIT^neg^ ILC2s (representative donor shown). (**G**) Heatmap showing scaled (row min-max) RPKM values for DEGs in the ILCP *vs*. KIT^neg^ ILC2 comparison (for clustering, see Methods). Selected genes are highlighted on the left. (**H**) mRNA expression levels of *RORC, LEF1*, and *CD40LG* in the indicated ILC subsets.

We first examined transcriptional differences between KIT^+^ and KIT^neg^ ILC2s (**Fig.2C**). Genes upregulated in KIT^neg^ ILC2s include known markers of type-2 lymphocyte biology (e.g., *GATA3-AS1*) but also DEGs associated with WNT and NFAT signaling (e.g., *DKK1*; **Fig.2C-D**). Interestingly, genes upregulated in KIT^+^ ILC2s were strongly linked to lymphocyte activation, cell migration, and lipid metabolism pathways (**Fig.2E**). These included genes encoding chemokine receptors such as *CCR6* and *CCR10*, which were previously shown to be expressed in KIT^+^ ILC2s^25^, but also the *MYC* master regulator of lymphocyte proliferation and metabolism^37^, the Oncostatin M (*OSM*) inflammatory cytokine, and co-stimulatory receptors GITR (*TNFRSF18*) and *CD99* (**Fig.2C, Supp.Fig.1A**). Importantly, flow cytometry analysis confirmed several of these transcriptional differences on the protein level (**Fig.2F**). Of note, only 24 genes were uniquely up- or downregulated in KIT^+^ ILC2s *vs*. both other ILC subsets **Supp.Fig.1B**), revealing that KIT^+^ ILC2s have very limited unique transcriptional features.

Next, we explored the transcriptional phenotype of KIT^+^ ILC2s in the context of ILCP-to-KIT^neg^ ILC2 differentiation. Clustering of DEGs in ILCP *vs*. KIT^neg^ ILC2 revealed 8 distinct expression patterns that confer a relative positioning of KIT^+^ ILC2s towards multipotent precursors and fate-committed ILC2s (**Fig.2G**). A small number of genes captured in clusters R1/R5 showed the highest or lowest expression in KIT^+^ ILC2s and play roles in regulating cell activation and migration (**Supp.Fig.1C**; partially overlapping with DEGs in **Fig.2C**), exemplified by *LEF1* encoding a TF that promotes *MYC* transcription^38^ and the CD40L co-stimulatory surface molecule (encoded by *CD40LG*; **Fig.2H**).

Many DEGs exhibited similar profiles across both ILC2 subsets (i.e., clusters R2, R7). Genes expressed highest in both KIT^+^ and KIT^neg^ ILC2s were linked to lymphocyte activation, interleukin signaling, and lipid metabolism (cluster R2; **Supp.Fig.1C**). These included known ILC2-associated genes, including TFs such as *RORA, MAF*, and *BCL11B* that control ILC2 identity^39–43^, and surface markers such as *KLRG1* and *IL9R* – the latter encoding the receptor for IL-9, a critical signal for ILC2 activation and survival^44,45^. Genes with the lowest expression levels in both ILC2 subsets were associated with broad immune cell activation pathways, integrin-mediated signaling, and responses to cytokines (Cluster R7; **Supp.Fig.1C**). Examples are *IL23R* and *CD244*, which encode surface receptors implicated in regulating ILCP differentiation^46,47^, but also co-stimulatory molecules such as *CD2* and *CD7* (**Fig.2G**).

In contrast, clusters R4 and R8 consisted of genes expressed to similar levels in ILCPs and KIT^+^ ILC2s (**Fig.2G**). These included genes involved in metabolic processes, growth factor responses and regulation of differentiation, including the *NR1D1* gene linked to mouse ILCP localization^48^ (**Fig.2G; Supp.Fig.1C**). *RORC*, encoding the master TF underlying ILC3 identity, showed similar robust expression levels in ILCPs and KIT^+^ ILC2s (**Fig.2H**). KIT^+^ ILC2s displayed intermediate expression levels for other clusters associated with cell activation and differentiation (**Fig.2G; Supp.Fig.1C**), involving genes important for both ILC2 (cluster R3; e.g. *IL13, CRLF2* (encoding the TSLP receptor), *RUNX1*) and ILCP (cluster R6: e.g. *TCF7, TOX, NCR1, ID2*) function.

Together, these observations reveal that KIT^+^ ILC2s display transcriptional characteristics of both KIT^neg^ ILC2s and ILCPs – akin to a developmentally ‘intermediate’ cell state. However, KIT^+^ ILC2s also upregulate a small yet unique transcriptional module that involves genes associated with cell activation, migration, and metabolism.

### IL-2 family cytokines and STAT5 induce a KIT^+^ ILC2-specific transcriptional program

TGFβ can induce expression of genes associated with a KIT^+^ ILC2 phenotype, including *RORC* and *CCR6*^25^. However, whether common γ-chain and alarmin cytokines – the canonical signals to which ILC2s respond – control the KIT^+^ ILC2-specific transcriptional program is unclear. To test this, we first exposed KIT^neg^ ILC2s to IL-2/IL-7 and observed a progressive increase in KIT^+^ cells (**Fig.3A**). Other genes significantly upregulated in KIT^+^ *vs*. KIT^neg^ ILC2s (as defined in **Fig.2C, Supp.Fig.1A**) also showed induction at the protein level (CD99, CCR10; **Fig.3A**). RNA-Seq of the total ILC2 population exposed to IL-2/IL-7 for 7 days showed robust and broad induction of the 137 genes that were significantly upregulated in KIT^+^ ILC2s compared to KIT^neg^ ILC2s (see **Fig.2C**; hereafter called the ‘KIT^+^ ILC2 gene signature’) (**Fig.3B-C**). Immune Response Enrichment Analysis (IREA^49^) on the KIT^+^ ILC2 gene signature identified IL-7 cytokine responses as the top association (enrichment score = 7.63, *P* = 0.02148) (**Fig.3D**). Notably, addition of alarmins severely blunted the upregulation of KIT^+^ ILC2 signature genes and proteins (**Fig.3E-H, Supp.Fig.2**).

**Figure 3.**
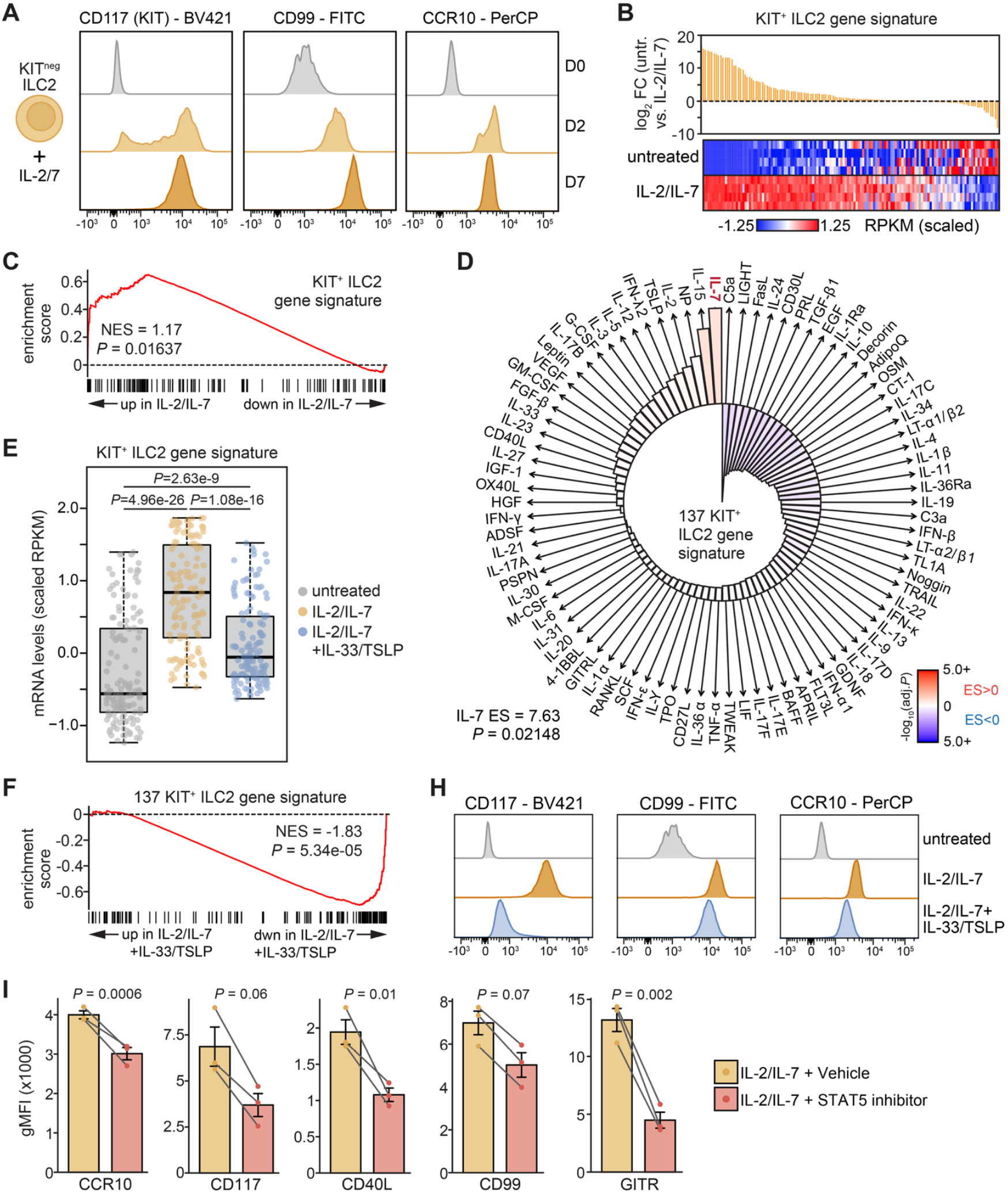
Common γ-chain and alarmin cytokines regulate the KIT^+^ ILC2-specific transcriptional program. (**A**) Flow cytometry analysis of KIT^neg^ ILC2s, untreated or cultured with IL-2/IL-7 for 2 or 7 days. (**B**) Expression (bulk RNA-Seq) of the KIT^+^ ILC2 gene signature (genes significantly upregulated in KIT^+^ ILC2s *vs*. KIT^neg^ ILC2s) on ILC2s untreated or cultured with IL-2/IL-7 for 7 days. Yellow bars show log_2_ fold changes (untreated *vs*. IL-2/IL-7). Heatmaps show scaled mRNA levels (z-score of RPKM) in individual biological replicates (n=4). (**C**) Gene set enrichment analysis (GSEA) of the KIT^+^ ILC2 gene signature in genes ranked on fold change between untreated and IL-2/IL-7-treated ILC2s. NES = normalized enrichment score. (**D**) Immune Response Enrichment Analysis (IREA) displaying enrichment of the KIT^+^ ILC2 gene signature among transcriptional programs induced in mouse ILCs by 86 different cytokines. Bar height denotes enrichment score (ES); color scale depicts *P*-values (only IL-7 and IL-15 showed *P*<0.05). (**E**) Box plot depicting averaged (n=4) scaled mRNA levels of the KIT^+^ ILC2 gene signature in untreated, IL-2/IL-7 and IL-2/IL-7 + IL-33/TSLP-treated ILC2 cultures (7 days). *P*-values: Wilcoxon rank sum test. (**F**) GSEA of the KIT^+^ ILC2 gene signature in genes ranked on fold change between IL-2/IL-7 and IL-2/IL-7 + IL-33/TSLP-treated ILC2s. (**H**) Flow cytometry analysis of sorted KIT^neg^ ILC2s, untreated or cultured for 7 days with IL-2/IL-7 or IL-2/IL-7 + IL-33/TSLP. (**I**) Geometric mean fluorescence intensity (gMFI) of indicated surface markers detected by flow cytometry on KIT^neg^ ILC2s cultured for 2 days with IL-2/IL-7 in the presence of DMSO (Vehicle) or 5 μM pimozide (i.e., STAT5 inhibitor). Bars denote mean values +/- SD obtained from 3 independent biological replicates. *P*-values: two-tailed T-test.

IL-2/IL-7 signaling results in STAT5 phosphorylation and its subsequent nuclear translocation to regulate gene expression^6^. To assess whether STAT5 activity is important for maintaining the KIT^+^ ILC2 phenotype, we exposed KIT^neg^ ILC2s to IL-2/IL-7 in the presence or absence of the STAT5 inhibitor pimozide. This revealed that STAT5 inhibition during a 48h exposure to IL-2/IL-7 reduced protein level induction of KIT^+^ ILC2 signature markers, including CCR10, KIT, CD40L, CD99 and GITR (encoded by *TNFRSF18*) (**Fig.3I**).

These experiments demonstrate that common γ-chain cytokines and downstream STAT5 activity induce the KIT^+^ ILC2 phenotype.

### KIT^+^ ILC2s harbor a plastic epigenomic landscape with progenitor-like features

We next analyzed the epigenomic landscapes of circulating ILCPs, KIT^+^ ILC2s, and KIT^neg^ ILC2s using ATAC-Seq. PCA of genome-wide chromatin accessibility positioned KIT^+^ ILC2s as an intermediate population along PC1 that separates ILCPs and KIT^neg^ ILC2s (**Fig.4A**). For example, the *GATA3* and TH2 cytokine loci (the latter encoding the type-2 cytokine genes *IL4, IL5*, and *IL13*) showed reduced chromatin accessibility in ILCPs (**Fig.4B**; **Supp.Fig.3A**). In KIT^+^ ILC2s, however, several gene-regulatory elements opened up, which further increased in accessibility in KIT^neg^ ILC2s (**Fig.4B**; gray shading). Conversely, the *TCF7* locus, encoding the TCF1 TF important for ILCP biology, showed more extensive accessibility in ILCPs and KIT^+^ ILC2s, whereas several of these regions were closed in KIT^neg^ ILC2s (**Fig.4B**; gray shading).

**Figure 4.**
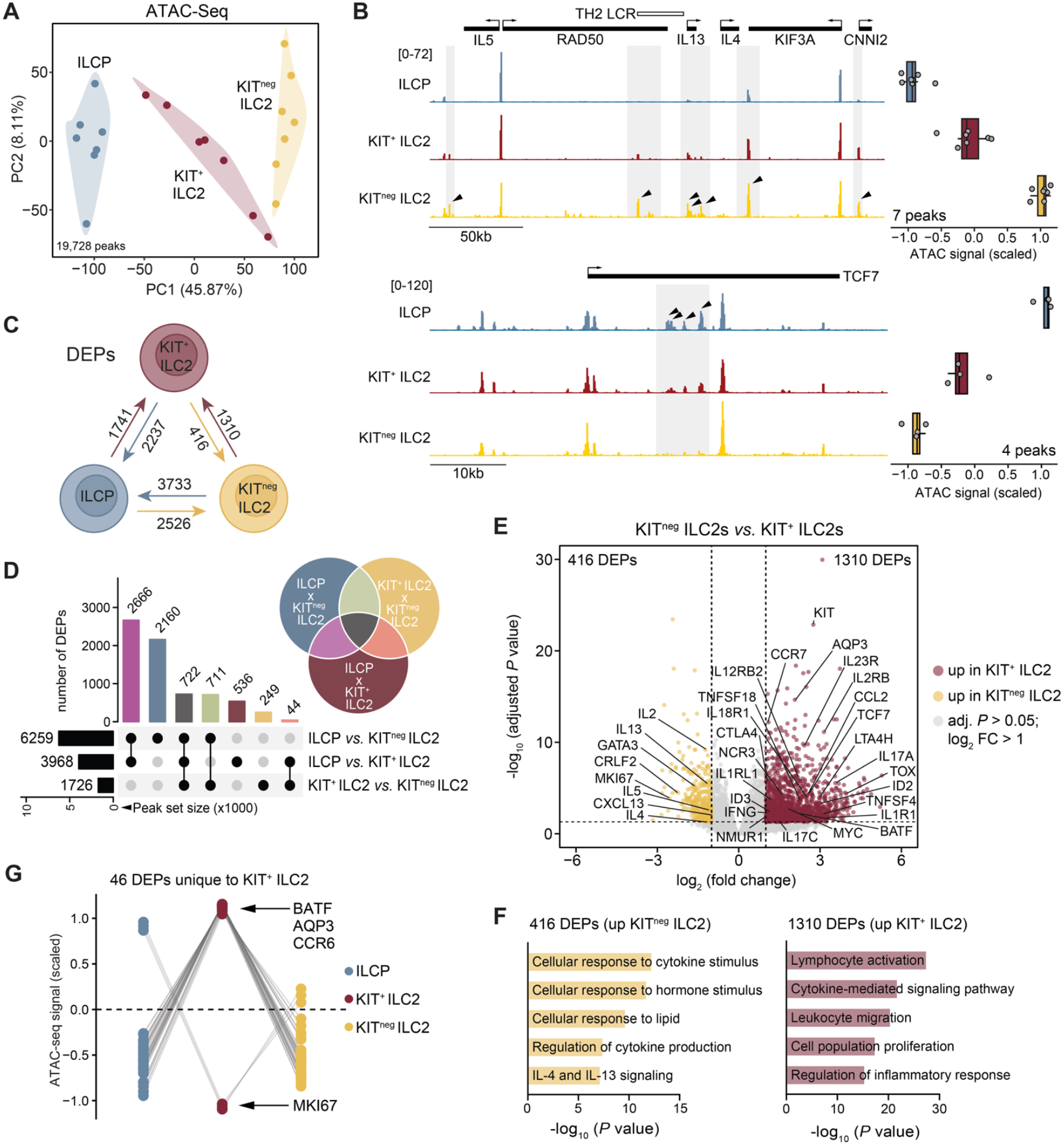
Distinct chromatin accessibility landscapes in KIT^+^ ILC2s. (**A**) PCA of normalized ATAC-Seq signals at all reproducible peaks detected in ILCPs, KIT^+^ ILC2s, and/or KIT^neg^ ILC2s (n=7 biological replicates per subset). (**B**) ATAC-Seq tracks (representative donor) at the TH2 cytokine locus (top) and *TCF7* locus (bottom). Grey shading highlights areas that contain differentially enriched peaks (DEPs; black arrows; log_2_ fold change > 1, adjusted *P* < 0.05). Box plots display scaled normalized counts per individual donor for all DEPs with increased accessibility in KIT^neg^ ILC2s (top) or ILCP (bottom). (**C**) Schematic overview of DEPs detected between ILCP, KIT^+^ ILC2s, and KIT^neg^ ILC2s. (**D**) Upset plot and Venn diagram visualizing the overlapping and unique DEPs among the indicated comparisons between ILC subsets. Color code matches across graphs. (**E**) Volcano plot showing DEPs between KIT^+^ ILC2s and KIT^neg^ ILC2s. Selected peak-associated genes are highlighted. (**F**) Bar graphs showing selected significantly enriched pathways for the genes associated with the DEPs depicted in panel E. (**G**) Scaled average normalized ATAC-Seq signals of 46 peaks uniquely up- or downregulated in KIT^+^ ILC2s compared to both ILCPs and KIT^neg^ ILC2s. Selected associated genes are indicated.

Genome-wide analysis detected the most differentially enriched peaks (DEPs) between ILCP and KIT^neg^ ILC2s (n=6259), and the fewest DEPs between KIT^+^ and KIT^neg^ ILC2s (n=1726) (**Fig.4C-D**). Comparing KIT^+^ and KIT^neg^ ILC2s, we observed that genes involved in type-2 cytokine signaling and cellular responses to hormones and lipids showed increased accessibility in KIT^neg^ ILC2s (**Fig.4E-F**). In KIT^+^ ILC2s, genes involved in lymphocyte activation, migration, and proliferation were more open (**Fig.4E-F**) – including many genes also detected in our transcriptome comparison (e.g., *MYC, OSM, CCR6*; 48 of the 137 upregulated DEGs). Still, the vast majority (96%) of epigenomically more accessible genes did not exhibit significantly increased steady-state mRNA levels in KIT^+^ ILC2s as compared to KIT^neg^ ILC2s (**Supp.Fig.3B**), indicative of broad epigenomic priming in KIT^+^ ILC2s that may facilitate future transcriptional upregulation. Notably, and similar to our RNA-seq results, KIT^+^ ILC2s displayed few (n=46) unique chromatin features (**Fig.4G**). These included DEPs associated with the KIT^+^ ILC2 marker gene *CCR6, AQP3* (linked to lymphocyte migration^50^), *BATF* – encoding a key regulator of ILC2 and ILC3 effector functions^51,52^ – and proliferation-associated marker *MKI67* (**Fig.4G; Supp.Fig.3C**). We next focused on DEPs between ILCPs and KIT^neg^ ILC2s and clustered them based on signal strength in KIT^+^ ILC2s (**Fig.5A**). Compared to findings at the transcriptome level (**Fig.2G**), a much larger fraction of DEPs displayed intermediate chromatin accessibility levels in KIT^+^ ILC2s – as reflected when comparing both PC1 scores (**Fig.5B**). DEPs in clusters A2-A4 were most accessible in KIT^neg^ ILC2s and exhibited various degrees of intermediate signal strength in KIT^+^ ILC2s (**Fig.5A**). Genes linked to these DEP clusters were associated with lymphocyte activation, (type-2) cytokine production, wound healing and cell adhesion, encoding canonical mediators of ILC2 function such as *RORA, BCL11B, GATA3* and *IL13* (**Fig.5C**). Clusters A6-A8 contained DEPs with highest ATAC-Seq signals in ILCPs and with varying intermediate levels in KIT^+^ ILC2s (**Fig.5A**). Here, DEPs localized near an overlapping set of genes that included ILCP-associated genes like *TCF7, TOX, ID2* and *CCR7*. Overall, genes linked to clusters A6-A8 were involved in diverse pathways, including leukocyte activation, stem cell differentiation, and NK cell-mediated immunity (**Fig.5C**). Notably, genes near DEPs from both opposing sets of clusters (i.e., A2-A4 *vs*. A6-A8) showed strong associations with cell activation and cytokine signaling pathways (**Supp.Fig.4A**), despite low general overlap (14.8%) in gene identity. However, a small set of genes was associated with multiple DEPs assigned to various opposing clusters (**Supp.Fig.4B**), indicating extensive and complex chromatin remodeling at a selective number of loci during human ILC2 differentiation. These loci harbor genes with key roles in ILC(2) biology (e.g., *GATA3, ID2, BACH2*) and lymphocyte activation (e.g., the *CD28*-*CTLA4*-*ICOS* locus, *MYC*) (**Supp.Fig.4C**).

**Figure 5.**
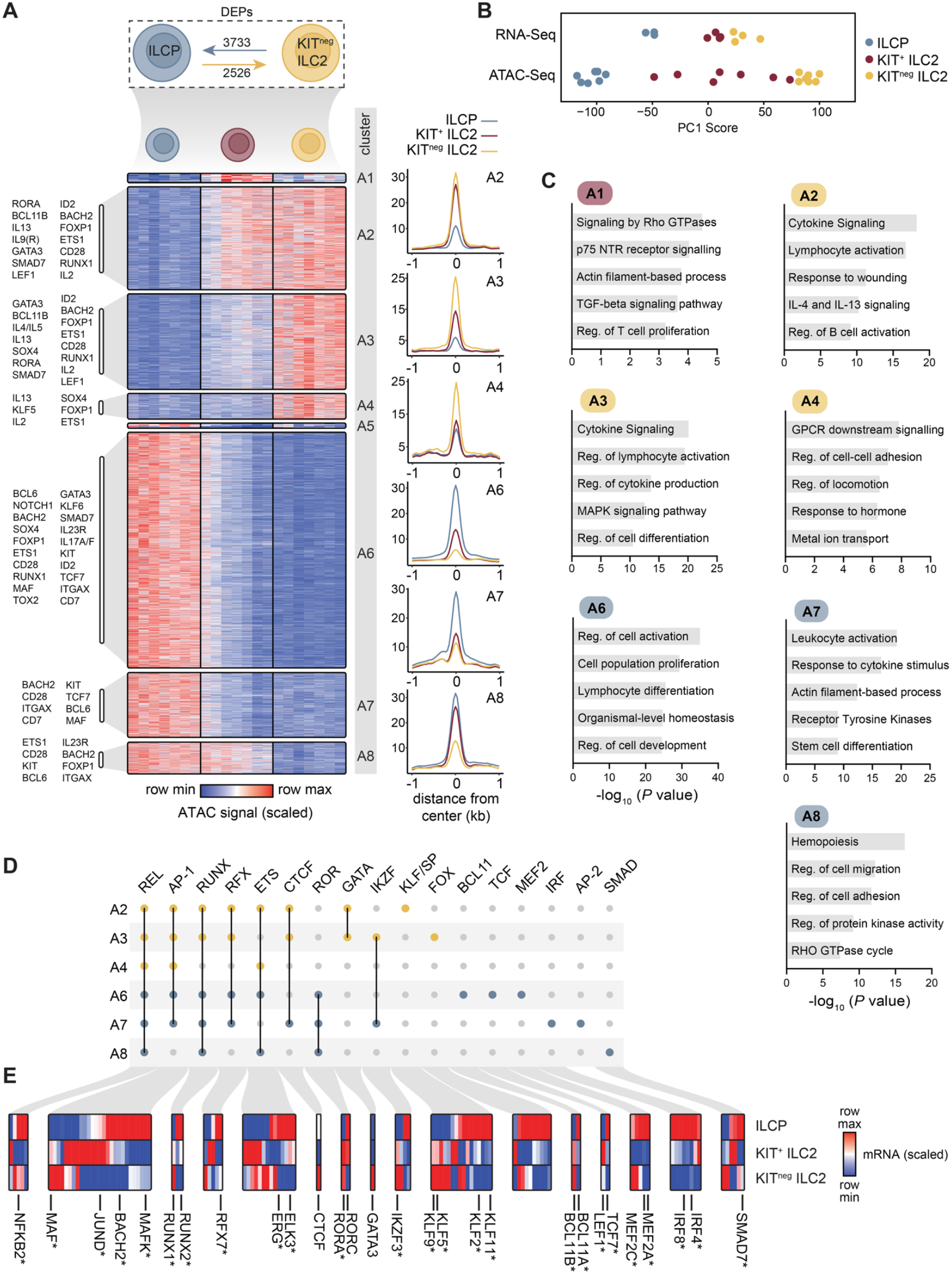
KIT^+^ ILC2s carry a broadly permissive epigenome with progenitor-like features. (**A**) Heatmap showing scaled (row min-max) normalized ATAC-Seq signals for all differentially enriched peaks (DEPs) in ILCPs *vs*. KIT^neg^ ILC2s (supervised clustering, see Methods). Selected genes associated with DEP clusters are indicated (left). Adjacent line graphs (right) display averaged normalized ATAC-seq signals across all peaks within the indicated clusters. (**B**) Plot showing PC1 values from PCAs performed on RNA-Seq (Fig.1C) or ATAC-Seq (Fig.4A) data. Each dot represents an individual sample, coloured by ILC subset. (**C**) Bar graphs showing selected significantly enriched pathways for the genes associated with the DEPs depicted in panel A. Reg. = Regulation. (**D**) Upset-style plot showing an overview of significantly enriched (see Methods) motifs per TF family in the indicated DEP clusters. (**E**) Heatmap displaying scaled (row min-max) mRNA levels of motif-associated TF genes (* indicates log_2_ fold change < 1 & adjusted *P*-value < 0.05).

To identify TFs that shape the intermediate epigenomic landscape of KIT^+^ ILC2s, we performed TF motif enrichment analysis on DEPs from clusters A2-4 and A6-8, and integrated results with our transcriptome data. Both peak sets (e.g., A2-4 & A6-8) were similarly enriched for REL (NF-κB), RFX, ETS and IKZF motifs (**Fig.5D, Supp.Table 2**), with differential gene expression observed for several associated TFs (e.g., *ERG, IKZF3*) (**Fig.5E**). ILCP-associated peaks (clusters A6-8) were more strongly enriched for RUNX, ROR, BCL11, TCF and IRF motifs (**Fig.5D, Supp.Table 2**), with concomitant increased expression of linked TF genes *RUNX2/3, RORC, BCL11A, TCF7* and *IRF4/8* in ILCPs or both KIT^+^ ILC subsets (**Fig.5E, Fig.2I, Supp.Fig.4D**). Interestingly, TCF7 can contribute to epigenetic priming of developmentally regulated genes in mouse ILCPs^53^. Intermediate peaks highest in KIT^neg^ ILC2s (clusters A2-4) were highly enriched in AP-1, GATA, KLF/SP and FOX family motifs (**Fig.5D, Supp.Table 2**), again accompanied by increased mRNA levels of type-2 lineage-associated TFs *MAF* (AP-1) and *GATA3* as well as several *KLF* genes (**Fig.5E, Supp.Fig.4D**). AP-1 family motifs also showed enrichment in clusters A6-7, along with elevated expression of *MAFK, JUND* and *BACH2* the latter displaying intermediate expression levels in KIT^+^ ILC2s (**Fig.5E, Supp.Fig.4D**). Despite the key role of STAT5 in driving the KIT^+^ ILC2 phenotype (**Fig.3I**), we did not observe enrichment of its motif in any peak set.

These observations reveal that KIT^+^ ILC2s carry a plastic epigenome that maintains progenitor-like features and may broadly prime them for future effector functions – beyond canonical ILC2-related activities. Opposing sets of TFs appear to maintain this ‘hybrid’ epigenomic landscape.

### KIT^+^ ILC2s maintain an epigenomically primed ILC3 program

Previous work has shown that KIT^+^ ILC2s more potently produce IL-17 as compared to KIT^neg^ ILC2s when exposed to type-3 inducing signals such as IL-23^25,26^, although the underlying mechanisms remain unclear. We hypothesized that epigenomic priming (i.e., accessible chromatin but none or very low transcription) of ILC3-associated loci in KIT^+^ ILC2s underlies their elevated plasticity towards an ILC3 phenotype. Indeed, ATAC-Seq analysis revealed highly accessible chromatin at the *IL23R* and *IL17* loci in resting KIT^+^ ILC2s, despite little to no transcription detected (**Fig.6A**). In contrast, KIT^neg^ ILC2s showed inaccessible chromatin, readily explaining the difference in IL-17 activation kinetics observed between both ILC2 subsets. We next identified 903 additional loci with a similar pattern of epigenomic priming in KIT^+^ ILC2s compared to KIT^neg^ ILC2s (see Methods). Most associated genes were not expressed in any ILC subset (∼80%, e.g., *IL17A*) or only in ILCPs (∼20%, e.g., *IL23R*) (**Fig.6B, Supp.Table 3**). Genes primed in KIT^+^ ILC2s were involved in cytokine signaling and leukocyte differentiation, including *IL23R, IL1RL1* (encoding the IL-33 receptor), *TNFSF18* (encoding the ligand for GITR), *IL12RB2* (encoding the IL-12 receptor), *IFNG*, and *IL17A* (**Fig.6B-C**; **Supp.Fig.4E**).

**Figure 6.**
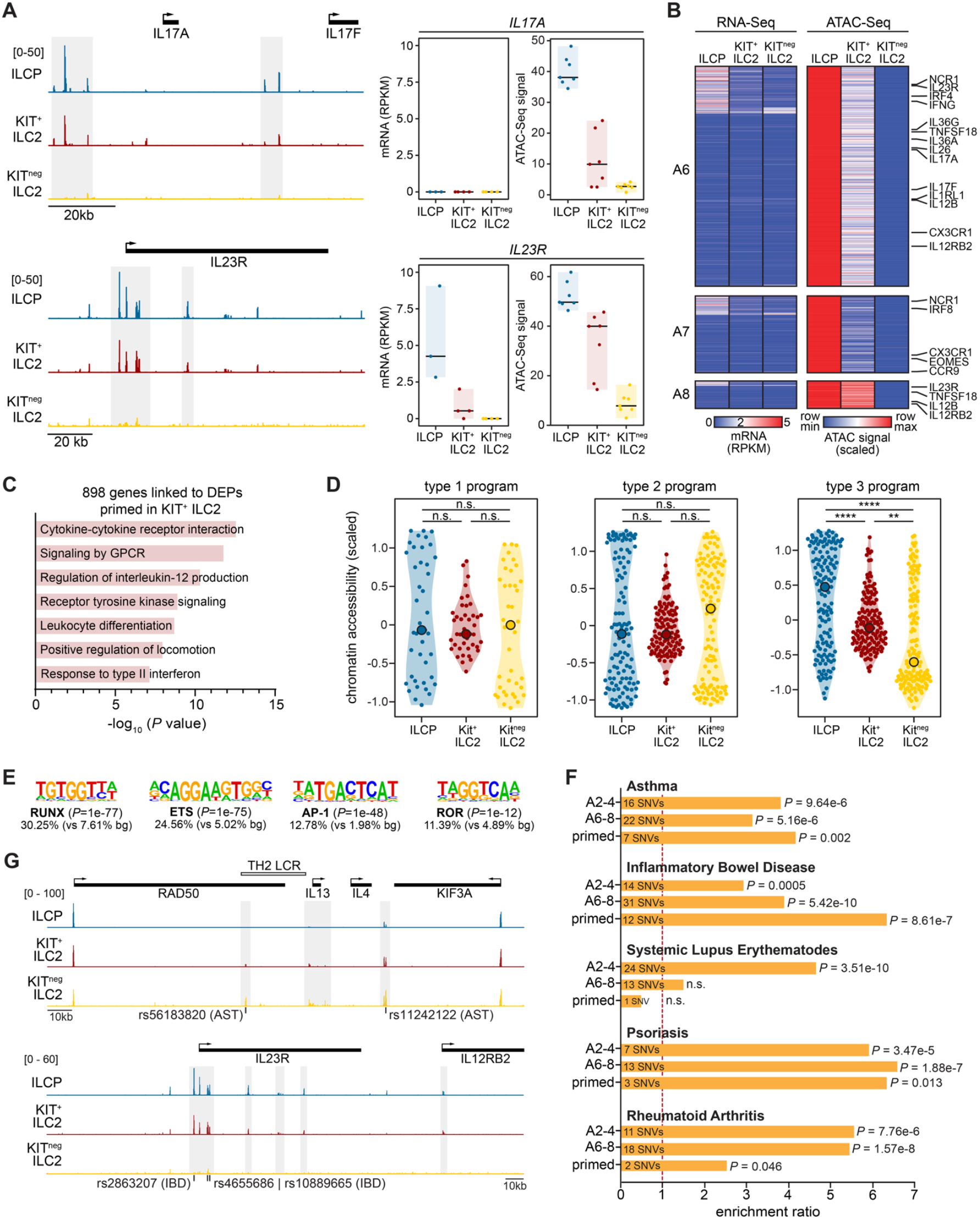
Epigenomic priming underlies enhanced KIT^+^ ILC2 plasticity and is linked to immune disease susceptibility. (**A**) Genome browser tracks (representative donor) showing ATAC-Seq profiles at the *IL17* and *IL23R* loci. Gray shading indicates DEPs between ILCP and KIT^neg^ ILC2 that appear primed in KIT^+^ ILC2. Box plots show transcription (RNA-Seq, left) and chromatin accessibility (ATAC-Seq, right - average signals at primed DEPs) per donor. (**B**) Heatmaps of loci (n=903) that are primed in KIT^+^ ILC2, i.e., low/no gene expression but substantial accessibility detected. Peaks were selected from intermediate clusters A6 (n=643), A7 (n=195) and A8 (n=65); see Fig.5A for original cluster definitions. RNA-Seq data are shown on the left (absolute RPKM counts averaged across donors), ATAC-Seq data on the right (scaled normalized count values averaged across donors for each DEP). (**C**) Bar graphs showing selected significantly (*P*<0.05) enriched pathways for the genes associated with the DEPs depicted in panel B. (**D**) Violin plots showing relative chromatin accessibility in ILCP, KIT^+^ ILC2, and KIT^neg^ ILC2 at loci associated with ILC1, ILC2, and ILC3 effector function or identity (literature-curated, see **Supp.Table 3**). For each gene, the peak with the maximum difference between ILCP and KIT^neg^ ILC2 was shown. (**E**) Top TF motifs enriched in primed regions. (**F**) Enrichment ratios and *P*-values (Fisher’s exact test) of genetic risk variants associated with asthma and indicated autoimmune diseases in peaks from clusters A2-4, clusters A6-8, and primed peaks. The number of overlapping single nucleotide variants (SNVs) is indicated. (**G**) Examples of disease-associated SNVs co-localizing with regulatory elements primed in KIT^+^ ILC2.

Using literature-curated type-1, type-2, and type-3 immunity-associated gene programs (**Supp.Table 4**), we observed overall increased accessibility at type-3 immunity-associated genes in ILCPs and KIT^+^ ILC2s as compared to KIT^neg^ ILC2s (**Fig.6D**). Various type-1 or type-2 immunity-associated genes were also primed in KIT^+^ ILC2s, although on average signals were more comparable between groups (**Fig.6D**). Indeed, binding motifs for RORγt (ROR) – a known key TF of type-3 lymphocyte identity – was enriched in primed peaks, alongside motifs for RUNX TFs known to be important for mouse ILC3 and ILC1 (but not ILC2) development^54^ (**Fig.6E**). *RORC* and *RUNX1-3* were expressed in KIT^+^ ILC2s, indicating that they may control priming at these sites (**Supp.Fig.4D**). We also observed motif enrichment and transcriptional activity of ETS (e.g., *ETS1*) and AP-1 (e.g., *JUNB*) factors (**Fig.6E, Supp.Fig.4D**), which are critical for ILC2 function^55,56^.

In summary, our analyses support epigenomic priming of inflammatory genes in KIT^+^ ILC2s as a mechanism underlying the high capacity of these cells for phenotypic plasticity towards an ILC3.

### Epigenomic priming in KIT^+^ ILC2s is linked to genetic susceptibility for immune diseases

IL-17 producing KIT^+^ ILC2s have been implicated in severe asthma and psoriasis^25,28^. A major risk factor for such diseases is common genetic variation, which can affect disease-relevant cellular phenotypes through modulating transcriptional regulation^57^. To investigate whether disease-associated genetic variants could impact KIT^+^ ILC2 biology, we integrated single nucleotide variants (SNVs) linked to asthma and several autoimmune disorders with our ATAC-Seq analyses. Overall, asthma and autoimmunity SNVs were enriched in both sets of intermediate peaks (i.e., A2-4 and A6-A8; see **Fig.5A**) and in the subset of 903 primed peaks (**Fig.6F**). Only systemic lupus erythematosus (SLE)-associated variants resided specifically in clusters A2-4. For asthma, intersecting SNVs localized near *GATA3, IL4* and *IL13* (clusters A2-4; **Fig.6G**) as well as *ID2, IL1R1* and *TNFSF18* (clusters A6-8) (**Supp.Table 5**). Genes linked to intersecting autoimmune disease-associated SNVs included *IL12RB2* and *IL23R* (**Fig.6G**), encoding receptors for cytokines with key roles in autoimmunity^58^.

Hence, epigenomic control of transcription in human ILC2s and their precursors – including priming of regulatory elements in KIT^+^ ILC2s – is linked to the genetic susceptibility for developing chronic inflammatory disease.

### Single-cell multi-omics reveals a developmentally intermediate epigenome in KIT^+^ ILC2s

To assess whether our findings can be extrapolated to the level of individual cells, we re-analyzed previously generated single-cell RNA-seq (scRNAseq) and scATACseq data from peripheral blood ILCs^35^. Falquet *et al*. identified two transcriptionally distinct populations of ILC2s labeled ‘ILC2a’ and ‘ILC2b’. Genes differentially expressed between ILC2a and ILC2b (n=58 DEGs) showed similar expression differences in our KIT^neg^ *vs*. KIT^+^ ILC2 transcriptome analysis (**Fig.7A**). In addition, our clusters of DEGs (**Fig.2G**) and DEPs (**Fig.5A**) showed comparable transcriptional and chromatin accessibility dynamics in the ILCP, ILC2a and ILC2b cells identified by Falquet *et al*. (**Fig.6B-C, Supp.Fig.5A**). We conclude that the ILC2a population is similar to our KIT^neg^ ILC2 population, whereas ILC2b cells very likely represent KIT^+^ ILC2s.

**Figure 7.**
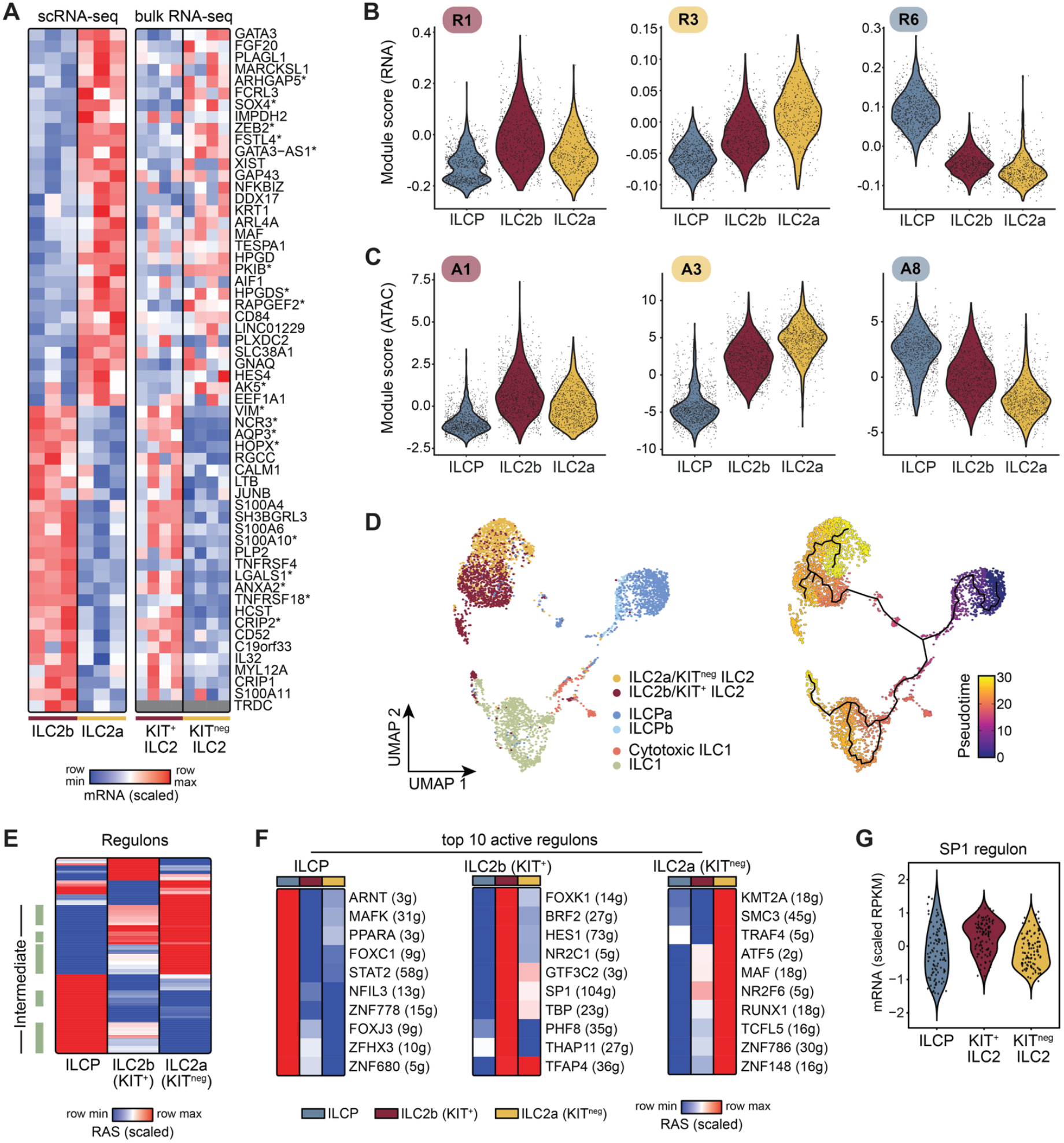
Single-cell transcriptome and epigenome analysis of human blood ILCs. (**A**) Scaled expression levels of DEGs from an ILC2a *vs*. ILC2b comparison^35^ (left heatmap) and a KIT^+^ ILC2 *vs*. KIT^neg^ ILC2 comparison (right heatmap). *indicates adjusted *P*<0.05 in bulk RNA-seq; grey color = no expression. (**B-C**) Violin plots showing the distribution of single-cell RNA-Seq (panel B) or ATAC-Seq (panel C) module scores for individual cells (dots). Scores reflect a combined value of all genes/peaks within the indicated bulk RNA-Seq (Fig.2G) or ATAC-Seq (Fig.5A) clusters. (**D**) Left: scATACseq data visualised by UMAP clustering (annotated by Falquet et al.). Right: Output of pseudo-time analysis – starting from the ILCPa cluster – superimposed on the left UMAP. (**E**) Heatmap showing scaled regulon activity scores (RAS) of all active regulons (rows) in the indicated ILC subsets. Regulons with intermediate activity scores in KIT^+^ ILC2s are labelled by green rectangles. (**F**) The 10 most differentially enriched regulons per ILC subset are indicated (adjusted *P*<0.05 in at least one subset). (**G**) Average bulk mRNA expression (scaled RPKM values) of the 104 genes within the SP1 regulon for each ILC subset.

We next performed a pseudotime analysis using the scATACseq dataset, using ILCPs as a starting point. This revealed a trajectory from ILCPs to ILC2a/KIT^neg^ ILC2s, with ILC2b/KIT^+^ ILC2s as an intermediate population (**Fig.7D**). A similar path was reported by Falquet *et al*. using their scRNAseq dataset^35^. These findings indicate that the KIT^+^ ILC2 phenotype may represent an intermediate stage during human ILCP-to-ILC2 differentiation.

To further dissect the transcriptional regulators that control ILCP-to-ILC2 differentiation, we used SCENIC to define TF-driven ‘regulons’ and measure their activity in each ILC subset. Regulons are modules of genes linked to a TF through a combination of co-expression and promoter motif enrichment^59^. Similar to our bulk RNA/ATAC analyses, ILC2b/KIT^+^ ILC2s frequently showed intermediate regulon activity scores, with few regulons showing predominant activation in ILC2b/KIT^+^ ILC2s (**Fig.7E**). ILCPs showed increased activity of the NFIL3 regulon, including the *KIT, LYL1*, and *CHD1* genes that have been linked to a progenitor phenotype^60,61^ (**Supp.Table 6)**. For ILC2a/KIT^neg^ ILC2s, MAF and RUNX1 regulons were highly active. Finally, ILC2b/KIT^+^ ILC2s showed high activity of SP1 and FOXK1 regulons (**Fig.7F, Supp.Fig.5B**), which include *NCR3* that encodes the NKp30 receptor important for ILC2 activation^62^, and the TF-encoding genes *IRF8* and *JUNB*. Indeed, SP1 regulon genes were on average also expressed at the highest levels in KIT^+^ ILC2s in our bulk RNA-Seq (**Fig.7G**).

Together, our analyses nominate KIT^+^ ILC2s as a plausible differentiation intermediate during ILCP-to-ILC2 differentiation, which terminates in a KIT^neg^ mature state. Moreover, we identified potential new regulators of human ILC2 differentiation and the KIT^+^ ILC2 phenotype.

## DISCUSSION

Our genome-wide analyses demonstrate that KIT^+^ ILC2s exhibit a bona fide type-2 lineage identity yet maintain a hybrid character marked by expression and chromatin accessibility of genes linked to precursor as well as type-2 effector states. Substantiated by single-cell analyses, our data position KIT^+^ ILC2s as a multi-potent intermediate stage, as KIT^+^ ILCPs differentiate into mature KIT^neg^ ILC2s. KIT^+^ ILC2s achieve enhanced phenotypic flexibility through a plastic precursor-like epigenome, poising genes for rapid activation – including *IL17* and *IL23R* that are key for plasticity towards an ILC3-like state. Moreover, these regions of primed chromatin are enriched for genetic variants linked to asthma and autoimmunity. Importantly, we show that common γ-chain cytokines IL-2 and IL-7 can induce a KIT^+^ phenotype in KIT^neg^ ILC2s through STAT5 activation, with important implications for therapeutic targeting of ILC2s populations (**Fig.8**).

**Figure 8.**
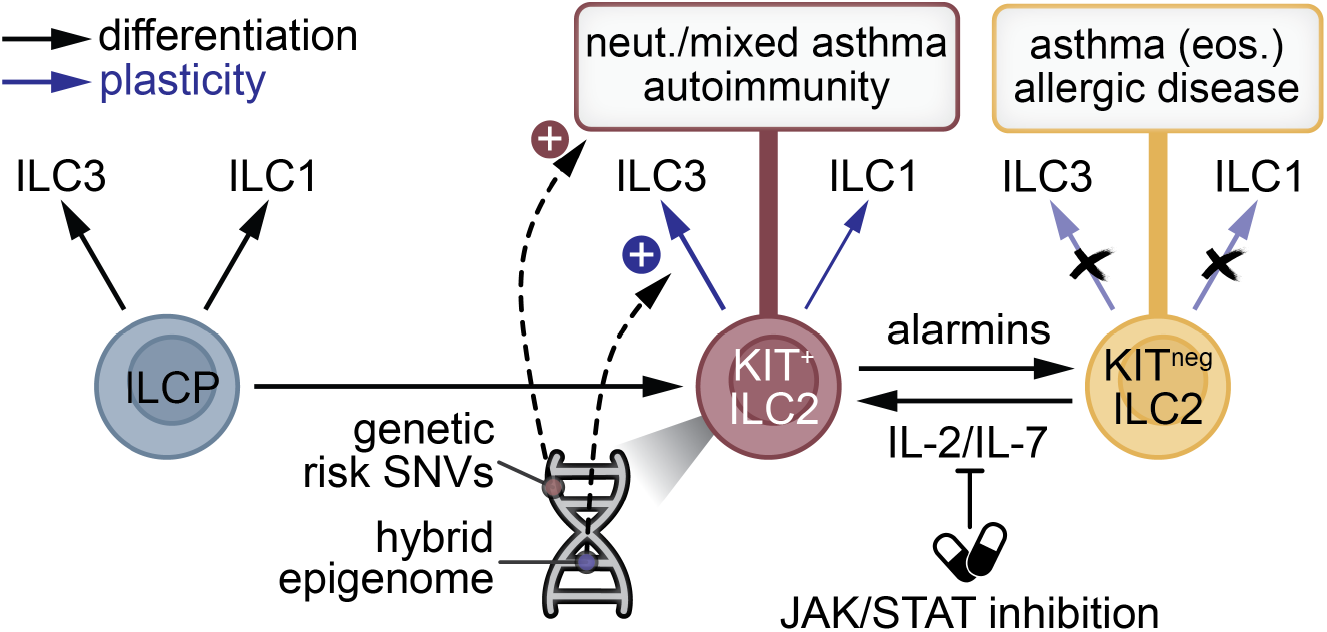
Schematic summarizing the main findings and implications of this study. See text for a detailed discussion. SNV: single nucleotide variant; neut.: neutrophilic; eos.: eosinophilic

Functional plasticity is widespread among the different ILC subsets^20,63^. The multipotent epigenome of KIT^+^ ILC2s we describe here readily explains the previously reported increased capacity of KIT^+^ ILC2s – in comparison with KIT^neg^ cells – to initiate IL-17 production under type-3 polarizing conditions^25,26^. While the molecular mechanisms that poise ILC2 epigenomes for functional plasticity are obscure, our data implicates regulatory regions, TFs, and signaling molecules in plasticity in KIT^+^ ILC2s. Among these, we observed TF family members with anti-correlated expression dynamics during ILCP-to-ILC2 differentiation, including *RORC*/*RORA, RUNX2*/*RUNX1*, and *BACH2*/*MAF*. We speculate that this may indicate antagonistic functions of TF family members at specific loci during human ILC2 differentiation, reminiscent of GATA TF switching described during hematopoietic development^64^. Both TFs in the above-mentioned pairs, linked to either ILCP- or KIT^neg^ ILC2-associated accessible chromatin, are expressed in KIT^+^ ILC2s, which may explain their hybrid multipotent epigenomic landscape and the complex chromatin dynamics observed at key cell identity loci (e.g., *GATA3, ID2, BACH2*).

Compared to ILCPs and KIT^neg^ ILC2s, we detected surprisingly small transcriptional (i.e. 24 genes) and epigenomic (i.e. 46 peaks) modules of specifically (more) active in KIT^+^ ILC2s, corroborating the notion that KIT^+^ ILC2s are epigenomically plastic but not a distinct ILC2 subset. Nevertheless, the KIT^+^ ILC2 gene signature does include genes encoding regulators of proliferation (*MYC*), co-stimulation (*CD40LG*), and cell migration (*CCR6, CCR10, AQP3*). Single-cell analyses further highlighted a set of TF-driven regulons specifically upregulated in KIT^+^ ILC2s, with SP1 emerging as a top hit that has not been implicated in ILC2 biology before. These observations suggest that KIT^+^ ILC2s may play functionally different physiological roles compared to KIT^neg^ ILC2s.

Our multi-omics analyses position KIT^+^ ILC2s as a potential intermediate state during differentiation of ILCPs into committed KIT^neg^ ILC2s. However, we also show that IL-2/IL-7 can induce KIT^neg^-to-KIT^+^ ILC2 conversion through STAT5 activation – akin to TGFβ^25^. Common γ-chain cytokines, in particular IL-7, are critical for ILC2 maintenance and function^65–67^. IL-7 levels in the tissue microenvironment, produced by for example epithelial and stromal cells^66,68,69^, may regulate switching between plastic KIT^+^ and committed KIT^neg^ states under steady-state conditions. Extrusion of tissue ILC2s^9^ recently exposed to IL-7 could explain the presence of KIT^+^ ILC2s in the circulation. As such, common γ-chain cytokines may shape an ILC2 population at barrier sites that offers a balanced capacity to drive canonical type-2 immunity and promote type-3 (or mixed) responses. Indeed, increased KIT^+^ ILC2s in sputum samples of severe asthmatics were specifically observed in patients with mixed eosinophilic/neutrophilic airway inflammation, together with elevated levels of type-3 inducing signals IL-1β and IL-18^28^. In agreement, our data revealed increased chromatin accessibility of the IL-18 receptor (*IL18R1*) and IL-1β receptor (*IL1R1*) loci in KIT^+^ as compared to KIT^neg^ ILC2s. Of note, we show that ILC2 plasticity seems to involve an alarmin-driven feedback loop, as IL-33 and TSLP suppressed the capacity of IL-2/IL-7 to induce a KIT^+^ multi-potent state. This is supported by our previous observations that alarmins induce a CD45RO^+^ KIT^neg^ ILC2 state resistant to plasticity, instead representing a major type 2 cytokine producer linked to severe (eosinophilic) type-2 respiratory diseases^12^.

Integrating GWAS data with our epigenomics datasets revealed substantial enrichment of asthma- and autoimmune-associated genetic variants in epigenomically primed regions in KIT^+^ ILC2s. This indicates that genetic susceptibility to various chronic inflammatory diseases may, in part, be explained by effects on (KIT^+^) ILC2 biology, further emphasizing the relevance of focusing on these cells in future research. The key role of STAT5 activation downstream of IL-2/IL-7 in promoting human ILC2 plasticity also offers new therapeutic angles that may be exploited to suppress pathological ILC2 activity – particularly in the context of IL-17-driven diseases. Compounds that effectively inhibit STAT5 or its upstream activator JAK3 exist^70,71^, and JAK3 inhibition was effective at suppressing lung ILC2 activation *in vivo*, including a subset of ILC2s producing both type-2 cytokines and IL-17^72^. Targeting these drugs to specific organs (e.g., inhalation into the lung^73^) could provide a viable strategy to specifically inhibit KIT^+^ ILC2 formation and activation in patients.

## ACKNOWLEDGEMENTS & FUNDING

The authors would like to thank members of the Pulmonary Medicine department at Erasmus MC for helpful discussions. This work received support from Lung Foundation Netherlands grants (4.1.18.226 to RWH; 4.2.19.041JO to RS and 6.1.25.069 to RS & RWH), a Dutch Research Council (ZonMw) Vidi grant (09150172010068 to RS), an Erasmus MC Fellowship (to RS), a Dutch Research Council (ZonMw) Open Competition grant (09120232310020 to RS & RWH), a Dutch Foundation for Asthma Prevention Grant (2022/007, to RS & RWH) and the EMBO Young Investigator Program (project number 5868, to RS).

