## Supplementary Data for "Common γ-chain cytokines induce an epigenomically plastic precursor-like KIT^+^ ILC2 state linked to immune disease susceptibility"

### AFFILIATIONS


### SUPPLEMENTARY MATERIALS

Supplementary Methods
Supplementary Figures S1-S5
Supplementary Tables 1-6

### METHODS

#### *Human blood samples and ILC isolation*

Buffy coats of healthy donors were provided by the Sanquin Blood Bank (Amsterdam, Netherlands; informed consent obtained by Sanquin). Peripheral blood mononuclear cells (PBMCs) were isolated by Ficoll-Paque Plus (Cytivia) density gradient centrifugation. To enrich for ILCs, PBMCs were depleted of T cells, B cells, monocytes, dendritic cells, platelets, and red blood cells by labeling with biotin-conjugated antibodies against CD3, CD4, CD19, CD14, CD16, CD36, and CD235αβ (**Supp.Table 1**), followed by magnetic-activated cell sorting using MojoSort Streptavidin Nanobeads (Biolegend) and LD columns (Mitenyi Biotec) following the manufacturer's instructions. After overnight incubation at 4°C in RPKM 1640 medium supplemented with 5% fetal calf serum (FCS), cells were stained with fluorescent antibodies (**Supp.Table 1**) for 30 min, followed by 15 min incubation with LIVE/DEAD Fixable Viability Dye eFluor506 (Thermo Fisher Scientific), both at 4°C in the dark. Then, ILCs were isolated using fluorescence-activated cell sorting on a FACSAria II (BDBiosciences) as Lineage-negative ($CD123^{neg}$, $CD14^{neg}$, $CD16^{neg}$, $CD19^{neg}$, $CD1a^{neg}$, $CD3^{neg}$, $CD34^{neg}$, $CD4^{neg}$, $CD5^{neg}$, $CD56^{neg}$, $CD94^{neg}$, $FC\varepsilon RI^{neg}$, $TCR\alpha/\beta^{neg}$, $TCR\gamma/\delta^{neg}$) $CD127^{+}CD3^{neg}$ cells. ILCPs were defined as $CD117^{+}CRTH2^{neg}$, $KIT^{+}$ ILC2 as $CD117^{+}CRTH2^{+,}$ and $KIT^{neg}$ ILC2 as $CD117^{neg}CRTH2^{+}$ (**Fig.1B**). For total ILC2 population isolation, $CD117^{+/-}CRTH2^{+}$ cells were sorted.

#### *In vitro culture of ILC2s and flow cytometry*

Sorted ILC populations were cultured for 2 or 7 days in Yssels's medium[75] supplemented with 1% normal human serum, IL-2 (10 U/ml, PeproTech), and IL-7 (20 ng/ml, PeproTech). In some cultures, IL-33, and TSLP (both 50 ng/mL, PeproTech) were added. To assess the role of STAT5 activity, $KIT^{+}$ ILC2s and $KIT^{neg}$ ILC2s were exposed to IL-2/IL-7 in the presence or absence of 5 μM Pimozide (TOCRIS Bioscience) or a vehicle control (0.5% DMSO). For flow cytometry readouts of both cultured and *ex vivo* samples, cells were stained extracellularly with fluorescent antibodies (**Supp. Table 1**) for 30 min at RT in the dark in PBS buffer. Fluorescence signals were measured on a FACSymphony A5 (BD Biosciences) and analyzed using FlowJo software (BD Biosciences).

#### *RNA isolation and bulk RNA sequencing*

Sorted cells were lysed in 350 μL RLT buffer (QIAGEN) supplemented with 1% β-mercapto-ethanol (Merck). RNA was extracted using the RNeasy Micro Kit (QIAGEN) according to the manufacturer's instructions. RNA-seq sequencing libraries were generated according to the Smart-seq2 method using the Nextera DNA Flex library prep kit (Illumina). Single 50 bp reads were generated using an Illumina HiSeq2500 sequencer. Illumina adapter sequences and poly-A stretches were trimmed from the reads using AdapterTrimmer[76], followed by read alignment to the hg38 human reference genome using HISAT2[77].

#### *Transcriptome analysis*

All aligned reads were parsed into tag directories using HOMER's[78] batchMakeTagDirectory.pl script. HOMER's analyzeRepeats.pl program was used to count reads at all RefSeq-annotated genes, counting only reads that mapped to exons. Read counts were normalized to reads per kilobase per million (RPKM). Differentially expressed genes (DEGs) were identified with DESeq2 through HOMER's getDiffExpression.pl script. Genes were considered differentially expressed if their expression differed between two ILC subsets (i.e., ILCP, $KIT^{+}$ ILC2, or $KIT^{neg}$ ILC2) or two culture conditions (i.e., untreated, IL-2/IL-7 or IL-2/IL-7/IL-33/TSLP) with a $\log_2$-fold change cut-off > 1 and an adjusted P-value < 0.05. In all downstream analyses, we only included genes that were expressed in at least one relevant group (RPKM > 1 in ≥50% of replicates). DEGs between ILCPs and $KIT^{neg}$ ILC2s were classified into clusters based on their relative expression in $KIT^{+}$ ILC2s. For each DEG, $\log_2$-fold-changes between ILCP *vs.* $KIT^{+}$ ILC2 and $KIT^{+}$ ILC2 *vs.* $KIT^{neg}$ ILC2 were used to assign genes into one of nine possible patterns: upregulated, downregulated, or equal in $KIT^{+}$ ILC2 relative to ILCP and $KIT^{neg}$ ILC2. Gene lists corresponding to each cluster were used for downstream analyses. One cluster did not contain any genes and was excluded from downstream analysis. PCA was performed on log2-transformed RPKMs of all genes that were expressed in at least one ILC subset using PCATools[79]. Pathway enrichment analysis was performed with Metascape[80]. Heatmaps were generated with Morpheus (https://software.broadinstitute.org/morpheus/). Gene set enrichment analysis (GSEA) was performed using clusterProfiler[81]: $Log_2$-fold changes from untreated *vs.* IL-2/IL-7-treated ILC2s or IL-2/IL-7 *vs.* IL-2/IL-7+IL-33/TSLP-treated cells were ranked and tested for enrichment of the 137 $KIT^+$ ILC2 signature genes (i.e., DEGs upregulated in $KIT^+$ ILC2s *vs.* $KIT^{neg}$ ILC2s). Immune Response Enrichment Analysis (IREA) was also performed on these 137 $KIT^+$ ILC2 signature genes[49].

#### *Bulk ATAC-Seq*

For ATAC-seq, cells were directly sorted in PBS + 10% FCS coated tubes containing 500 µL cold lysis buffer (0.3 M Sucrose, 10 mM Tris-HCl HCL pH 7.5, 60 mM KCL, 15 mM NaCl, 5 mM MgCl, 5 mM $MgCl_2$, 0.1 mM EGTA, 0.10% NP40, 0.15 mM Spermine, 0.5 mM Spermidine, 2 mM 6AA in MilliQ). Cells were centrifuged for 10 min at 500 x g at 4 °C, and the cell pellet was resuspended in 25 µL 2X reaction buffer (Nextera kit, Illumina) with 2.5 µL Nextera Tn5 transposase (Illumina) and 22.5 µL nuclease-free $H_2O$. The reaction mixture was incubated for 45 min at 37°C, while shaking gently at 500 rpm. Immediately after transposition, the suspension was purified using the MinElute PCR Purification kit (QIAGEN) according to the manufacturer's instructions. DNA fragments were amplified (13-15 cycles, depending on the number of sorted cells and DNA concentration) and Illumina sequence adapters were added by PCR: 1 cycle of 5 min 72ºC and 30 sec 98ºC, followed by 13-15 cycles of 10 sec 98ºC, 30 sec 63ºC, and 1 min 72ºC. Libraries were purified and fragments <100 base pairs or >1000 base pairs removed using AMPure XP beads (Beckman Coulter). Paired-end 50 bp reads were generated using an Illumina NextSeq2000 sequencer. Sequence reads were trimmed using AdapterTrimmer[76] and aligned to the hg38 human reference genome using HISAT2[77].

#### *Bulk ATAC-Seq data analysis*

Sequence reads were filtered for a maximum fragment length of 500 base pairs. Duplicated reads were removed using sambamba markdup --remove-duplicates[82]. Additionally, deepTools was used to remove reads mapped to blacklisted genomic regions and to shift reads for the Tn5 cutting site using alignmentSieve --ATACshift --blackListFileName[83,84]. Tag directories were created for all samples using HOMER's batchMakeTagDirectory.pl, with reads mapping to mitochondrial DNA and the Y chromosome removed[78]. Peaks were called using HOMER (findPeaks -region -localSize 75000 -size 75 -minDist 75). Peaks detected in at least four out of seven replicates of at least one ILC subset were considered reproducible and included for downstream analysis. HOMER's annotatePeaks.pl was used to determine raw read counts per reproducible peak (-size given -raw). To visualize the data with the IGV genome browser, raw bedGraphs (makeUCSCfile -raw - fsize 50e6) were generated with HOMER. GREAT was used with default association rule settings to assign each peak to up to two genes that it is most likely to regulate[85]. The average coverage at peak sets of interest was calculated with HOMER (annotatePeaks.pl hg38 -size 10000 -hist 40 -raw). Raw counts were normalized using scaling factors calculated with DESeq2[86] based on a common set of peaks that was reproducibly found in all three ILC subsets as previously described[87]. PCA was performed on log2-transformed DESeq2-normalized peak counts of all reproducible peaks using PCATools. Differentially enriched peaks (DEPs; $log_2$-fold change > 1, adjusted P-value < 0.05) were identified using DESeq2, controlling for batch in the design[86]. DEPs between ILCPs and $KIT^{neg}$ ILC2s were grouped into clusters based on their relative accessibility in $KIT^+$ ILC2s. For each DEP, $log_2$-fold-changes between ILCP *vs.* $KIT^+$ ILC2 and $KIT^+$ ILC2 *vs.* $KIT^{neg}$ ILC2 were used to assign genes into one of nine possible patterns: upregulated, downregulated, or equal in $KIT^+$ ILC2 relative to ILCP and $KIT^{neg}$ ILC2. Peak lists corresponding to each cluster were used for downstream analyses. However, the cluster representing peaks that were similar between KIT+ ILC2s and both ILCP and KITneg ILC2s was excluded from downstream analysis, since it contained only 1 peak. To identify epigenomic priming in KIT+ ILC2s, peaks from clusters A6-A8 were filtered based on their associated gene(s) not being expressed in KIT+ ILC2s. Epigenetic changes at genes associated with type-1, type-2, and type-3 associated gene signatures were captured by focusing on the mean difference at the gene-associated peaks that captured the maximum average difference between ILCP and KITneg ILC2. Pathway enrichment analysis was conducted using Metascape[80]. TF motif enrichment analysis was performed on differentially accessible regions using HOMER's findMotifsGenome command (-size 200 -mask -len 6,8,10,12 -S 20).

***Single nucleotide variant (SNV) enrichment analysis***

Immune disease-associated genetic variants were obtained from published genome-wide association studies[88–92]. Summary statistics files were parsed to the Functional Mapping and Annotation (FUMA) platform[93], which uses PLINK to impute missing genotypes from the 1000 Genomes reference panel[94]. Default settings were used. Genomic locations of the resulting SNVs were converted from GRCh37/hg19 to GRCh38/hg38 coordinates using the LiftOver tool of the UCSC Genome Browser. To determine enrichment of disease-associated SNVs within sets of ATAC-Seq peaks, we calculated whether the overlap between SNVs and peaks was greater than expected by chance given their sizes and the size of the hg38 reference genome using BEDTools' Fisher's exact test[95].

***Single-cell RNA-seq data analysis***

Publicly available single-cell RNA-seq data (GSE225169) were obtained as count matrices and were analyzed using the Seurat package (v5.0.1)[96]. Data processing, dimensionality reduction, clustering, and annotation of ILC subsets were performed following the workflow described by Falquet et al.[35]. In addition, normalized expression values for selected genes were extracted using the addModuleScore function in Seurat and visualized across the identified clusters of our bulk RNA-seq data.

***SCENIC analysis***

Gene regulatory network inference was performed on the processed single-cell RNA-seq dataset using pySCENIC[97]. The calculated regulon activity scores were compared for each ILC cluster (i.e., ILCP, ILC2a, ILC2b) against all other cells using Wilcoxon rank-sum tests with Benjamini-Hochberg correction for multiple testing. Regulons statistically significant for at least one cluster (adjusted $P < 0.05$) were retained. Within each ILC cluster, the regulons were ranked by $\log_2$-fold change, with the top ten most enriched selected for downstream analysis.

***Single-cell ATAC-seq data analysis***

Publicly available single-cell ATAC-seq data (GSE225169) were obtained as fragment files and analyzed using Signac (v1.12.9004), GenomicRanges (v1.54.1), EnsDb.Hsapiens.v86 (v2.99.0), Cicero (v1.3.9), and JASPAR2020 (v0.99.10), following an approach similar to that described by Falquet et al.[35]. Trajectory analysis was performed using Monocle3 (v1.3.4)[98], with the ILCPa cluster defined as the starting point. Normalized accessibility scores for selected peaks identified in our bulk ATAC-seq data (>50% overlap in genomic coordinates with single-cell ATAC-seq peaks) were calculated using the AddModuleScore function in Seurat.

***Data availability statement***

In order to comply with privacy legislation for data sharing purposes, the aligned RNA-Seq and ATAC-Seq BAM files generated for this study were anonymized using BAMboozle v0.5.0[74]. Anonymized BAM files and tables containing quality control metrics, raw and normalized transcript counts were deposited in the Zenodo repository (DOI: 10.5281/zenodo.17788534).

**Supplementary Figure 1** – Olsthoorn, Onrust et al.

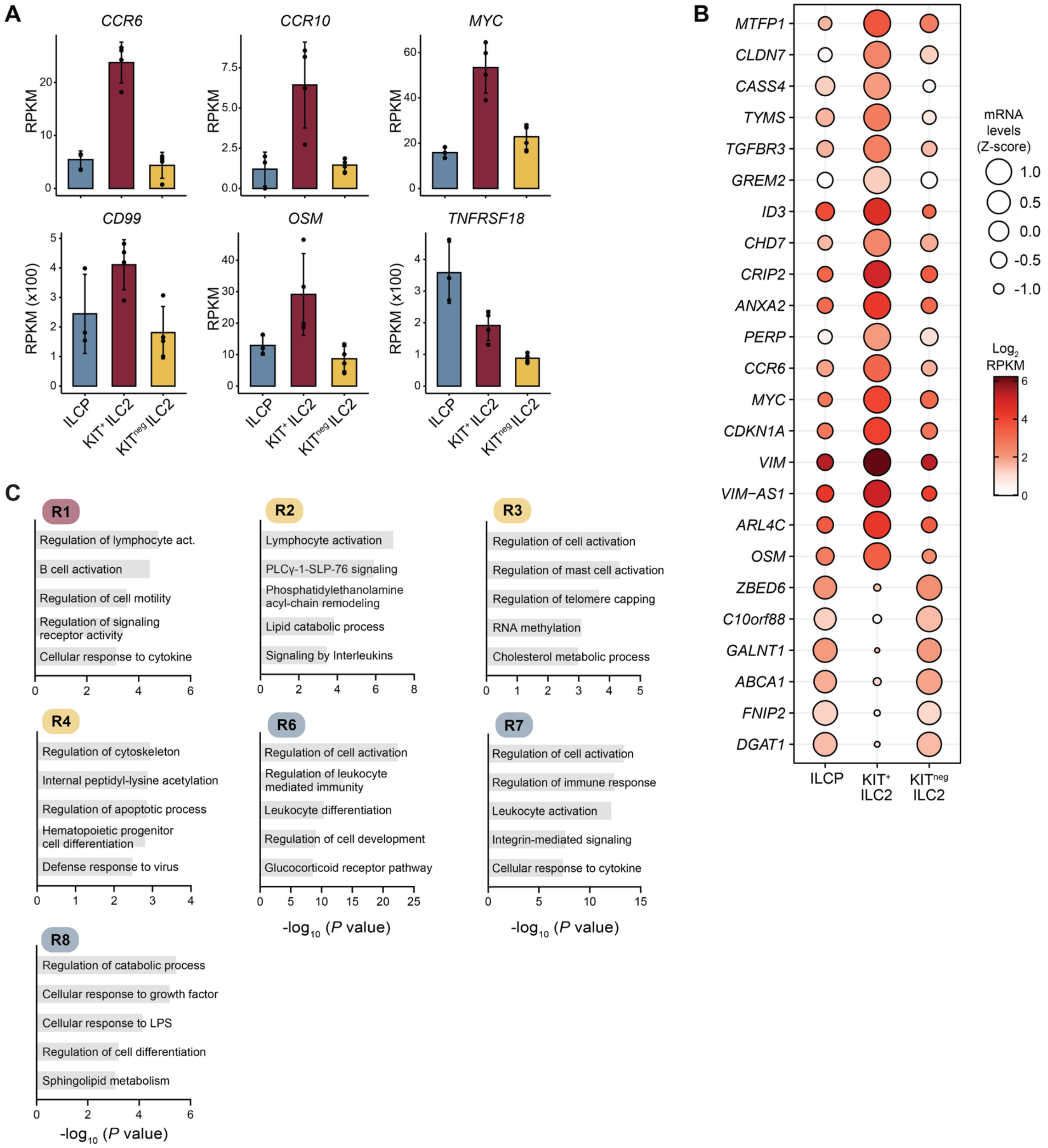


**Supplementary Figure 1.** (**A**) Bar graphs showing mRNA expression levels of *CCR6, CCR10, MYC, CD99, OSM,* and *TNFRSF18* in the indicated ILC subsets. Bars indicate mean values with error bars showing ±1 SD. (**B**) Balloon plot depicting mRNA expression levels in the indicated ILC subsets of the 24 DEGs that were uniquely up- and downregulated in KIT$^{+}$ ILC2s compared to both ILCPs and KIT$^{neg}$ ILC2s. Circle size indicates relative expression to the other subsets (z-score), while the color scale shows absolute expression levels ($log_2$ RPKM). (**C**) Bar graphs showing selected significantly enriched pathways for the clusters of genes depicted in Fig.2G.

**Supplementary Figure 2** – Olsthoorn, Onrust et al.

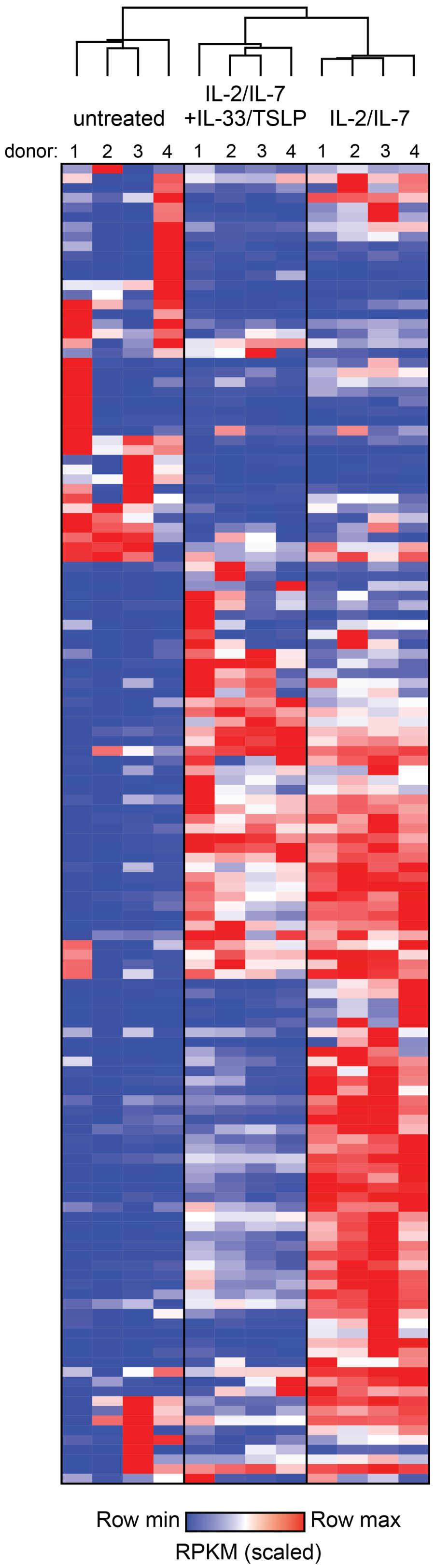


**Supplementary Figure 2.** Heatmap showing scaled (row min-max) RPKM values for the KIT$^{+}$ ILC2 gene signature in untreated ILC2s, ILC2s treated with IL-2/IL-7, and ILC2s treated with IL-2/IL-7 + IL-33/TSLP (n=4 biological replicates). Columns represent individual replicates hierarchically clustered (rows and columns).

**Supplementary Figure 3** – Olsthoorn, Onrust et al.

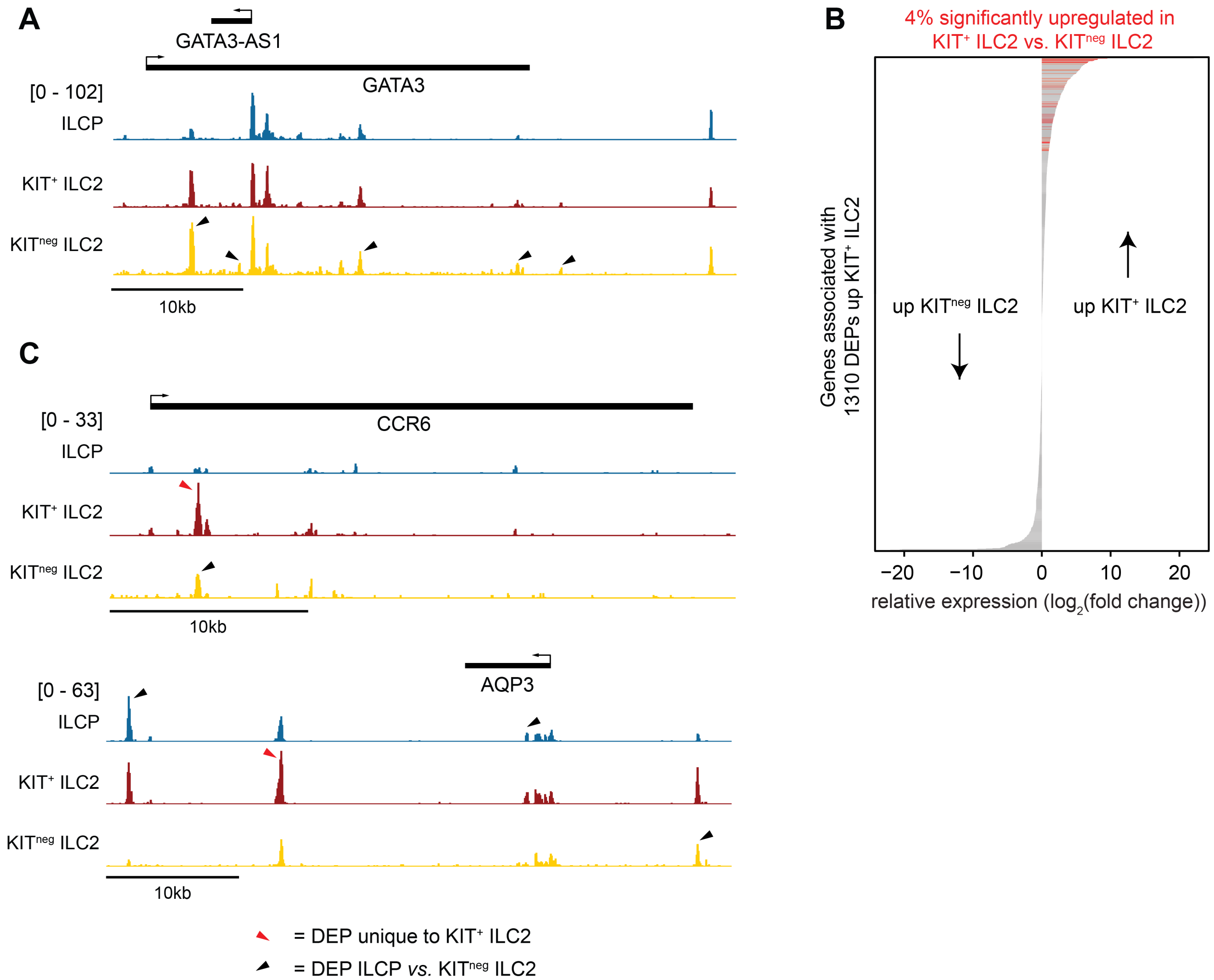


**Supplementary Figure 3. (A)** Representative ATAC-Seq track displaying chromatin accessibility at the *GATA3* locus. Black arrows indicate DEPs between ILCP and KIT$^{neg}$ ILC2. (**B**) Relative transcription of genes associated with 1310 DEPs that showed increased accessibility in KIT$^{+}$ ILC2s *vs.* KIT$^{neg}$ ILC2s. Plotted is the $\log_2$ fold change between KIT$^{+}$ *vs.* KIT$^{neg}$ ILC2s for each gene (x-axis) in descending order (y-axis). Significantly higher expressed genes in KIT$^{+}$ ILC2s compared to KIT$^{neg}$ ILC2s are indicated as red bars (log2 fold change > 1 & adjusted P-value < 0.05; 4% of all genes). (**C**) Representative ATAC-Seq tracks displaying chromatin accessibility at *CCR6* (top) and *AQP3* (bottom) loci. Red arrows highlight DEPs uniquely increased in KIT$^{+}$ ILC2s (see Fig.4D, orange set). Black arrows indicate DEPs between ILCP and KIT$^{neg}$ ILC2.

**Supplementary Figure 4** – Olsthoorn, Onrust et al.

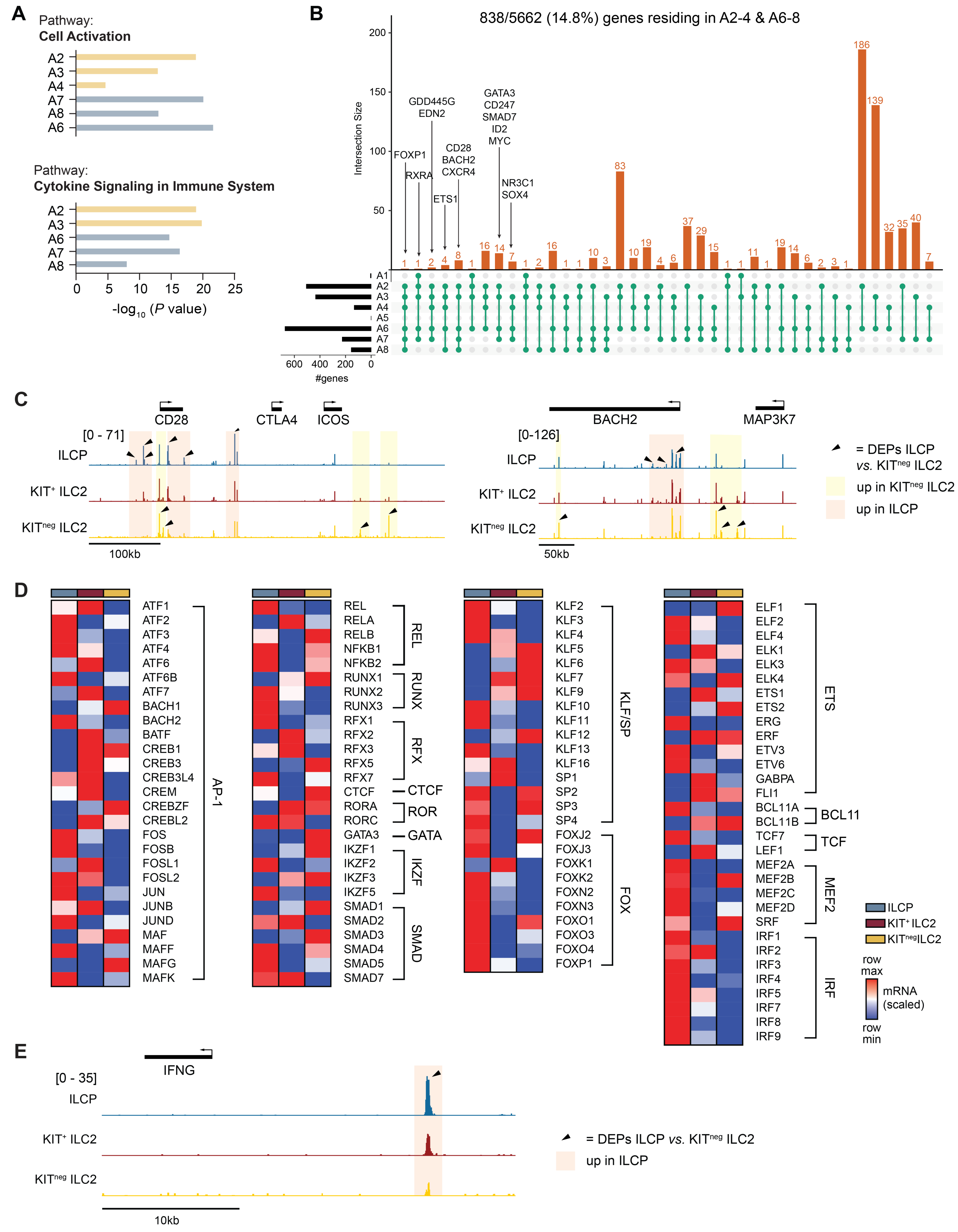

**Supplementary Figure 4.** (**A**) Bar graphs showing $-\log_{10}$ enrichment scores for 'cell activation' and 'cytokine signalling in immune system' pathways across the indicated ATAC-Seq clusters (shown in Fig.5A). Yellow bars indicate clusters containing peaks most accessible in $KIT^{neg}$ ILC2s; blue bars indicate clusters with peaks most accessible in ILCPs. (**B**) Upset plot visualizing the overlaps between genes associated with ATAC-seq peak clusters from Fig.5A. Only genes appearing at least once in clusters A2-4 and at least once in clusters A6-8 are shown (838/5662 genes). Selected genes with substantial overlap between opposing sets of clusters (i.e., A2-A4 *vs.* A6-A8) are highlighted. (**C**) Representative ATAC-Seq tracks displaying chromatin accessibility at *CD28-CTLA4-ICOS* (left) and *BACH2* (right) loci. Black arrows indicate DEPs between ILCP and $KIT^{neg}$ ILC2, shading colors denote direction of change. (**D**) Heatmaps showing average scaled (row min-max) RPKM values for all TFs related to the enriched TF family motifs as shown in Fig.5D. Columns represent ILCP, $KIT^{+}$ ILC2s and $KIT^{neg}$ ILC2s. (**E**) Representative ATAC-Seq tracks displaying chromatin accessibility at *IFNG* loci. Black arrows indicate DEPs between ILCP and $KIT^{neg}$ ILC2, shading colors denote direction of change.

**Supplementary Figure 5** – Olsthoorn, Onrust et al.

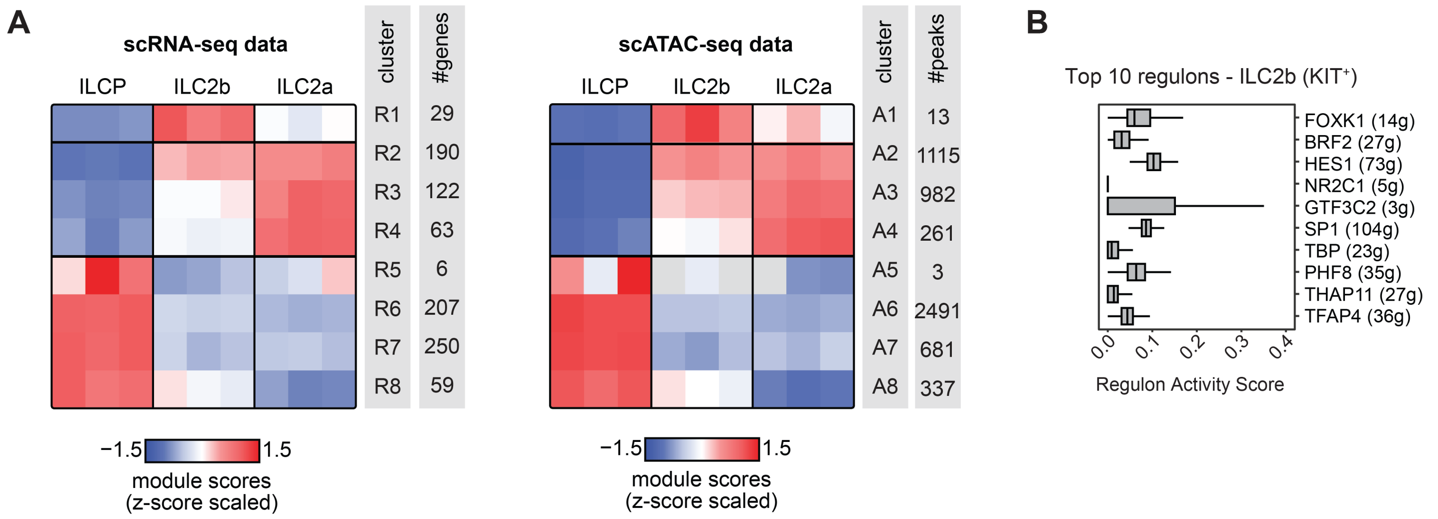


**Supplementary Figure 5.** (**A**) Heatmaps showing average scaled module scores of the indicated clusters as calculated in the single cell (sc)RNA-Seq (left) and scATAC-Seq (right) data. Scores reflect a combined value of all genes/peaks within the indicated bulk RNA-Seq (R clusters, Fig.2G) or ATAC-Seq (A clusters, Fig.5A) clusters. The number of genes or peaks in each cluster is indicated. (**B**) Boxplots depicting regulon activity scores of the top 10 most enriched regulons in ILC2b.

**Supplementary Table 1 – Antibody information**

| Target | Conjugate | Clone | Company | Identifier |
|---|---|---|---|---|
| CCR10 | PerCP | 1B5 | BD Biosciences | 564772 |
| CD117 (c-kit) | BV421 | 104D2 | Biolegend | 313216 |
| CD123 | FITC | 6H6 | Biolegend | 306014 |
| CD127 | APC | eBioRDR5 | Life Technologies | 17-1278-42 |
| CD14 | FITC | 61D3 | Life Technologies | 11-0149-42 |
| CD14 | Biotin | 61D3 | Life Technologies | 13-0149-82 |
| CD154 (CD40L) | BV650 | 24-31 | Biolegend | 310863 |
| CD16 | FITC | 3G8 | BD Biosciences | 555406 |
| CD16 | Biotin | eBioCB16 | Life Technologies | 13-0168-82 |
| CD19 | FITC | HIB19 | BD Biosciences | 555412 |
| CD19 | Biotin | HIB19 | Life Technologies | 13-0199-82 |
| CD196 (CCR6) | BV605 | G034E3 | Biolegend | 353420 |
| CD1a | FITC | HI149 | Biolegend | 300104 |
| CD235ab | Biotin | HIR2 | Biolegend | 306618 |
| CD294 (CRTH2) | PE/Cyanine7 | BM16 | Biolegend | 350118 |
| CD294 (CRTH2) | PE-Texas Red | BM16 | Biolegend | 350125 |
| CD3 | FITC | UCHT1 | Life Technologies | 11-0038-42 |
| CD3 | APC-Cy7 | SK7 | BD Biosciences | 557832 |
| CD3 | Biotin | UCHT1 | Life Technologies | 13-0038-82 |
| CD335 (NKp46, NCR1) | PE | 9E2 | Biolegend | 331907 |
| CD34 | FITC | 581 | Biolegend | 343504 |
| CD357 (GITR) | BV711 | 108-17 | Biolegend | 371211 |
| CD36 | Biotin | 5-271 | Biolegend | 336218 |
| CD4 | FITC | OKT4 | Biolegend | 317408 |
| CD4 | Biotin | OKT4 | Life Technologies | 13-0048-82 |
| CD45RA | PE-Texas Red | MEM-56 | Life Technologies | MHCD45RA17 |
| CD5 | FITC | UCHT2 | Biolegend | 300606 |
| CD56 (NCAM) | FITC | HCD56 | Biolegend | 318304 |
| CD94 | FITC | DX22 | Biolegend | 305504 |
| CD99 | FITC | hec2 | Biolegend | 398207 |
| FceR1α | FITC | AER-37 (CRA1) | Life Technologies | 11-5899-42 |
| KLRG1 | APC | 13F12F2 | Life Technologies | 17-9488-42 |
| TCR α/β | FITC | IP26 | Biolegend | 306706 |
| TCR γ/δ | FITC | B1 | Biolegend | 331208 |

### Supplementary Table 2 – Transcription factor motif enrichment per ATAC-Seq cluster

| A2 | | | A3 | | | A4 | | | A6 | | | A7 | | | A8 | | |
|---|---|---|---|---|---|---|---|---|---|---|---|---|---|---|---|---|---|
| *de novo* Motif | family | enrichment | *de novo* Motif | family | enrichment | *de novo* Motif | family | enrichment | *de novo* Motif | family | enrichment | *de novo* Motif | family | enrichment | *de novo* Motif | family | enrichment |
| ERG | ETS | P=1e-72 27.69% vs. 8.18% bg | REL | REL | P=1e-56 16.61% vs. 3.27% bg | NFkb-p65 | REL | P=1e-20 19.34% vs. 3.53% bg | RUNX | RUNX | P=1e-284 39.53% vs. 10.58% bg | IKZF1 | IKZF | P=1e-28 18.29% vs. 5.56% bg | GABPA | ETS | P=1e-57 34.60% vs. 4.69% bg |
| Jun-AP1 | AP-1 | P=1e-40 13.79% vs. 3.54% bg | FOSL2:JUNB | AP-1 | P=1e-56 17.78% vs. 3.86% bg | CREB1 | AP-1 | P=1e-15 24.28% vs. 7.35% bg | ERG | ETS | P=1e-247 34.50% vs. 9.02% bg | AP-1 | AP-1 | P=1e-28 12.80% vs. 2.92% bg | RUNX-AML | RUNX | P=1e-32 37.01% vs. 10.39% bg |
| REL | REL | P=1e-39 12.98% vs. 3.19% bg | Gata1 | GATA | P=1e-43 30.01% vs. 12.15% bg | BATF3 | AP-1 | P=1e-15 14.40% vs. 2.66% bg | Fra1 | AP-1 | P=1e-160 17.75% vs. 3.42% bg | NFkB-p65 | REL | P=1e-24 11.59% vs. 2.76% bg | Smad2::Smad3 | SMAD | P=1e-25 19.72% vs. 3.54% bg |
| Atf2 | AP-1 | P=1e-28 20.79% vs. 9.05% bg | IKZF1 | IKZF | P=1e-36 16.72% vs. 5.02% bg | ETV1 | ETS | P=1e-13 7.00% vs. 0.55% bg | REL | REL | P=1e-92 16.75% vs. 5.05% bg | RUNX2 | RUNX | P=1e-23 20.73% vs. 8.01% bg | RORg | ROR | P=1e-17 9.00% vs. 0.88% bg |
| Gata4 | GATA | P=1e-28 15.62% vs. 5.72% bg | JUN | AP-1 | P=1e-32 20.05% vs. 7.41% bg | | | | Bcl11a | BCL11 | P=1e-59 42.33% vs. 26.47% bg | THRb | ROR | P=1e-20 49.24% vs. 31.70% bg | NFkB-p50p52 | REL | P=1e-16 5.19% vs. 0.19% bg |
| RUNX1 | RUNX | P=1e-27 19.47% vs. 8.39% bg | RUNX1 | RUNX | P=1e-29 8.82% vs. 1.77% bg | | | | Tcf7 | TCF | P=1e-51 16.00% vs. 6.78% bg | BORIS | CTCF | P=1e-18 10.52% vs. 2.97% bg | | | |
| Atf1 | AP-1 | P=1e-24 16.63% vs. 6.96% bg | Rfx5 | RFX | P=1e-26 10.08% vs. 2.52% bg | | | | RORA | ROR | P=1e-31 13.25% vs. 6.38% bg | RFX1 | RFX | P=1e-17 8.38% vs. 2.06% bg | | | |
| CTCF | CTCF | P=1e-20 12.37% vs. 4.73% bg | CTCF | CTCF | P=1e-16 2.63% vs. 0.22% bg | | | | Mef2d | MEF2 | P=1e-27 4.33% vs. 1.15% bg | IRF4 | IRF | P=1e-15 31.86% vs. 18.65% bg | | | |
| Sp1 | KLF/SP | P=1e-19 21.70% vs. 11.45% bg | Atf1 | AP-1 | P=1e-15 4.12% vs. 0.71% bg | | | | PU.1 | ETS | P=1e-23 4.68% vs. 1.50% bg | TFAP2C | AP-2 | P=1e-14 17.68% vs. 8.09% bg | | | |
| RFX1 | RFX | P=1e-16 2.43% vs. 0.23% bg | Foxd3 | FOX | P=1e-15 5.15% vs. 1.15% bg | | | | RFX7 | RFX | P=1e-21 6.47% vs. 2.65% bg | Ptf1a | - | P=1e-14 2.29% vs. 0.11% bg | | | |

*bg: background*

**Supplementary Table 3 – Primed genes & associated ATAC-Seq peaks**

*See excel file*

**Supplementary Table 4 – Type 1, 2 and 3 gene signatures**

*See excel file*

**Supplementary Table 5 – ATAC-Seq cluster peak intersect with GWAS SNVs**

*See excel file*

**Supplementary Table 6 – Regulons detected by SCENIC analysis**

*See excel file*
